# A modular molecular framework for quickly estimating the binding affinity of the spike protein of SARS-CoV-2 variants for ACE2, in presence of mutations at the spike receptor binding domain

**DOI:** 10.1101/2021.05.26.445422

**Authors:** Vincenzo Tragni, Francesca Preziusi, Luna Laera, Angelo Onofrio, Simona Todisco, Mariateresa Volpicella, Anna De Grassi, Ciro Leonardo Pierri

## Abstract

The rapid spread of new SARS-CoV-2 variants needs the development of rapid tools for predicting the affinity of the mutated proteins responsible for the infection, i.e., the SARS-CoV-2 spike protein, for the human ACE2 receptor, aiming to understand if a variant can be more efficient in invading host cells. Here we show how our computational pipeline, previously used for studying SARS-CoV-2 spike receptor binding domain (RBD)/ACE2 interactions and pre-/post-fusion conformational changes, can be used for predicting binding affinities of the human ACE2 receptor for the spike protein RBD of the characterized infectious variants of concern/interest B.1.1.7-UK (carrying the mutations N501Y, S494P, E484K at the RBD), P.1-Japan/Brazil (RBD mutations: K417N/T, E484K, N501Y), B.1.351-South Africa (RBD mutations: K417N, E484K, N501Y), B.1.427/B.1.429-California (RBD mutations: L452R), the B.1.141 variant (RBD mutations: N439K), and the recent B.1.617.1-India (RBD mutations: L452R; E484Q) and the B.1.620 (RBD mutations: S477N; E484K). Furthermore, we searched for ACE2 structurally related proteins that might be involved in interactions with the SARS-CoV-2 spike protein, in those tissues showing low ACE2 expression, revealing two new proteins, THOP1 and NLN, deserving to be investigated for their possible inclusion in the group of host-cell entry factors responsible for host-cell SARS-CoV-2 invasion and immunity response.

## Introduction

The spread of the new SARS-CoV-2 variants makes necessary the development of new tools able to predict binding affinities of the SARS-CoV-2 spike protein, with the human dipeptidyl carboxydipeptidase angiotensin converting enzyme II (ACE2) receptor [1–3]. This knowledge might be crucial for evaluating the increased affinity of the spike protein of the new variants of concern (VoC) for the human ACE2 receptor. Indeed, a greater spike/ACE2 affinity may result in a loss of efficiency of vaccines based on a specific spike elder sequence, i.e., the hCoV.19Wuhan.WIV04.2019 [4, 5]. In addition, the obtained information concerning more dangerous (in terms of binding affinity) VoC may help in planning future antibodies [1, 6] and/or rounds of vaccination by using new vaccines based on a cocktail of the newly identified spike sequences (in the form of mRNA [4, 5] or proteins/subdomains [7–9]) showing high affinities for the human host-cell receptors.

In this research context our laboratory provided a modular molecular framework for investigating the molecular mechanism that allows SARS-CoV-2 entry in human host cells through SARS-CoV-2 spike receptor binding domain (RBD)/ACE2 interactions [1, 2]. However, the crystallized structure of RBD/ACE2 protein complex from 6vw1.pdb, used in our previous work [1], as the only one available at the time (March 2020), consisted of a chimeric SARS-CoV-2 spike RBD [10] and showed 22 missense mutations and a deletion of 3 amino acids at the RBD, with respect to the amino acid sequence of Wuhan SARS-CoV-2 spike protein (as reported in the YP_009724390.1 sequence, namely the hCoV.19Wuhan.WIV04.2019 sequence, https://www.gisaid.org/hcov19-variants/). Thus, we re-estimated the binding interactions between the human ACE2 receptor and SARS-CoV-2 spike RBD, both reported in the new available crystallized structure of the Wuhan SARS-CoV-2 spike protein/ACE2 (6m0j.pdb) protein complex [11].

It is known that mutations occurring at the RBD are the most dangerous due to their ability in increasing binding affinity for ACE2, favoring virus entry into human host-cells and antibody escape [12–15]. Thus, aiming to investigate the effect of amino acid replacement at the SARS-CoV-2 spike RBD, we built a 3D comparative model for each spike variant showing an amino acid replacement at the RBD positions K417; N439; L452, E484; S477; S494; N501, as highlighted in the SARS-CoV-2 VoC B.1.1.7-United Kingdom (UK, N501Y, S494P, E484K), P.1-Japan/Brazil (K417N/T, E484K, N501Y), B.1.351-South Africa (SA, K417N, E484K, N501Y), B.1.427/B.1.429-California (L452R), the B1.141 variant (N439K) and the recent B.1.617.1-India (L452R; E484Q) and the B.1.620 (showing two mutations at the spike RBD, namely S477N; E484K). Notably, all the cited VoC genomes have been sequenced in the last year and are responsible for the current pandemics situation [15–21].

Finally, given the low expression of ACE2 observed in some tissues highly exposed to SARS-CoV-2, and the acquired knowledge about a set of possible human host-cell entry factors [22–25], we used our computational pipeline for predicting ACE2 structurally related receptors, aiming to identify other putative SARS-CoV-2 entry sites in human host cells, based on a folding recognition approach, more than on sequence/functional comparative analysis.

## Materials and methods

### Comparative 3D modelling of SARS-CoV-2 spike RBD and the investigated RBD mutants interacting with the human ACE2 receptor

Starting from the atomic coordinates of the SARS-CoV-2 spike RBD protein domain extracted from 6m0j.pdb (according to the YP_009724390.1 sequence), we built the 3D comparative models of the investigated RBD mutants with specific reference to the single mutants N501Y, E484K/Q, N439K, K417N/T, L452R, S477N, S494P, the double mutants L452R-E484Q; and S477N-E484K, and the triple mutants N501Y-E484K-K417N; N501Y-E484K-S494P, N501Y-E484K-K417T by using the mutagenesis tool implemented in SwissPDBViewer [26].

In order to obtain a pose of the 3D protein complex of SARS-CoV-2 spike RBD mutants interacting with ACE2, the built 3D comparative models of the mutants were superimposed to the 3D protein complex consisting of the Wuhan SARS-CoV-2 spike RBD interacting with ACE2, available under the 6m0j.pdb protein data bank (PDB) entry.

All the generated 3D all-atom models were energetically minimized using the Yasara Minimization server and residues packing was checked and repaired, where necessary, according to the FOLDX repair function [27]. PyMol [28] (https://www.pymol.org) was then used for examining (by manual inspection) the obtained 3D structure models, and for checking the correct packing of local secondary structures.

### Crystal structure sampling of possible ACE2 structural related alternative receptors via folding recognition and multiple sequence alignments (MSA)

ACE2 structure related proteins were sampled by using the folding recognition methods implemented in pGenThreader [29] and I-Tasser [30]. With this aim, the amino acid sequence of ACE2 (NP_001358344.1) was used as query sequence for running pGenThreader (http://bioinf.cs.ucl.ac.uk/psipred/) and I-Tasser (https://zhanglab.ccmb.med.umich.edu/I-TASSER/) to screen the Protein Data Bank (PDB), searching for ACE2 structurally related crystallized proteins [1, 29–35].

The crystallized structures of the proteins sampled by pGenThreader and I-Tasser analysis were structurally aligned with the 3D coordinates of ACE2 available under the PDB_ID 6m0j.pdb. For obtaining the structural alignment we used the “super” command available in Pymol [28], which is able to structurally align also proteins with a lower percentage of identical residues, due to its ability in providing a sequence-independent structure-based pairwise alignment [1, 31, 33, 34].

The sequences of the investigated crystallized ACE2 structurally related proteins were aligned using ClustaW [36] and optimized by visual inspection based on the structural alignment obtained by PyMOL [28].

### Preparation of 3D complex protein models hosting the Wuhan SARS-CoV-2 spike RBD in complex with the highlighted ACE2 structurally related receptors

In order to obtain the most likely 3D protein complexes of SARS-CoV-2 spike RBD interacting with the highlighted ACE2 structurally related receptors “dipeptidyl carboxydipeptidase angiotensin I converting enzyme 1 (ACE, https://www.ncbi.nlm.nih.gov/gene/1636)”, “thymet oligopeptidase 1 (THOP1, https://www.ncbi.nlm.nih.gov/gene/7064)”, and “neurolysin peptidase (NLN, https://www.ncbi.nlm.nih.gov/gene/57486)”, the sampled crystallized structures of ACE, THOP1 and NLN were superimposed on the structure of the human ACE2 receptor within 6m0j.pdb, crystallized in complex with the SARS-CoV-2 spike RBD.

The obtained 3D coordinates of SARS-CoV-2 spike RBD (according to the YP_009724390.1 sequence as obtained from 6m0j.pdb) in complex with ACE, THOP1, and NLN were each saved in a new PDB file.

The variation in the number of interactions (H-bonds, ionic and aromatic interactions) at the SARS-CoV-2 spike RBD/ACE2 interface in presence of the investigated amino acid replacements have been calculated by using the PIC webserver [37] and verified by manual inspection by using PyMOL.

### FoldX energy calculations

The 3D coordinates of the Wuhan SARS-CoV-2 spike RBD crystallized in complex with the human ACE2 receptor available under the PDB_ID 6m0j.pdb were used to estimate the binding affinity of the SARS-CoV-2 spike RBD for the human ACE2 by using the FoldX AnalyseComplex assay [38]. The calculated binding affinity of the two protein domains within the crystallized 6m0j.pdb was used as a reference value for the following comparative analyses [1].

Indeed, the FoldX AnalyseComplex assay was performed to determine the interaction energy between the investigated minimized protein complexes consisting of the SARS-COV-2 spike RBD mutants (N501Y, E484K/Q, N439K, K417N/T, L452R, S477N, S494P, the double mutants L452R-E484Q, S477N-E484K, and the triple mutants N501Y-E484K-K417N, N501Y-E484K-S494P and N501Y-E484K-K417T) and ACE2.

Furthermore, the FoldX AnalyseComplex assay was used for determining the binding affinity between SARS-CoV-2 spike RBD and the ACE2 structurally related receptors ACE, THOP1, and NLN, identified through pGenTHREADER and/or ITASSER analyses.

The way the FoldX AnalyseComplex operates is by unfolding the selected targets and determining the stability of the remaining molecules and then subtracting the sum of the individual energies from global energy [1]. More negative energies indicate a better binding, whereas positive energies indicate no binding [38, 39].

### GTEx expression map of human coronavirus entry factors

The GTEx portal [40] was screened to assess the gene expression levels of ACE2 structurally related receptors, i.e., ACE, THOP1 and NLN, and of many other genes proposed to play a crucial role in host-cell virus entry processes, i.e., the immunoglobulin basigin (BSG, also known as CD147, https://www.ncbi.nlm.nih.gov/gene/682), the interferon-induced membrane protein IFITM3 (known for being involved in the protection against several viruses, https://www.ncbi.nlm.nih.gov/gene/10410), the FURIN protease (https://www.ncbi.nlm.nih.gov/gene/5045), the lysosomal cysteine peptidases CTSB (https://www.ncbi.nlm.nih.gov/gene/1508) and CTSL (https://www.ncbi.nlm.nih.gov/gene/1514), the membrane alanyl aminopeptidase ANPEP (https://www.ncbi.nlm.nih.gov/gene/290), the transmembrane serine proteases TMPRSS2 (https://www.ncbi.nlm.nih.gov/gene/7113) and TMPRSS4 (https://www.ncbi.nlm.nih.gov/gene/56649), the dipeptidyl peptidase DPP4 (also known as CD26, https://www.ncbi.nlm.nih.gov/gene/1803), the glycan-binding receptors of the C-type lectin family (known for being involved in the recognition of several viruses and bacteria) CLEC4G (https://www.ncbi.nlm.nih.gov/gene/339390) and CLEC4M (https://www.ncbi.nlm.nih.gov/gene/10332), the lymphocyte antigen 6E LY6E (https://www.ncbi.nlm.nih.gov/gene/4061) [24, 25].

## RESULTS

### RBD amino acid replacements, number of interactions, local secondary structure perturbation, and interaction energy calculation

The analysis about SARS-CoV-2 VoC spread in the last year has highlighted the important role played by amino acid replacements occurring at the spike RBD in SARS-CoV-2 host-cell entry and virus infection [1–3, 12–15, 41–43]. Here we report about the employment of a computational pipeline for the estimation of binding affinity and interactions between SARS-CoV-2 spike RBDs and ACE2 in presence of mutations occurring at seven positions of the spike RBD, as observed in seven VoC (Fig. 1), chosen among the best characterized variants (https://www.ecdc.europa.eu/en/covid-19/variants-concern).

**Figure 1.**
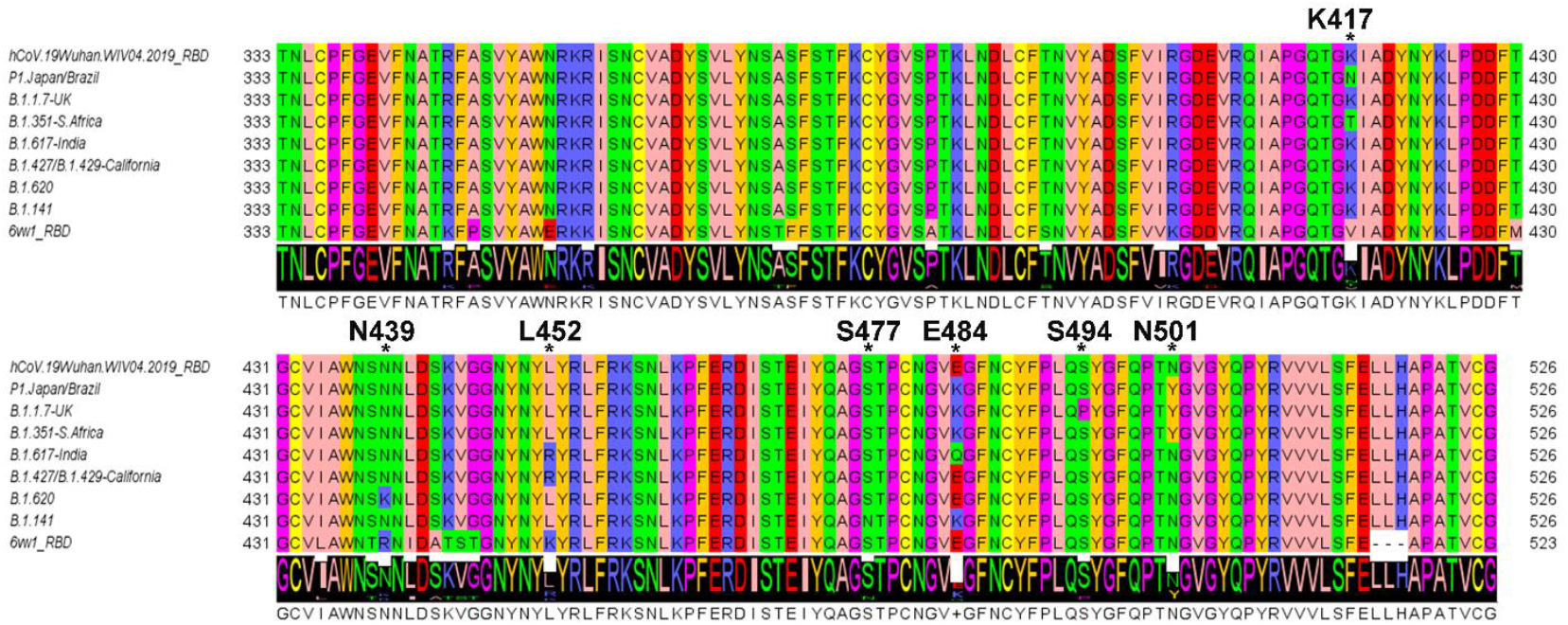
Multiple sequence alignment (MSA) of SARS-CoV-2 spike RBD highlighted from the sequenced VoC. MSA of the SARS-CoV-2 spike RBD from 6m0j.pdb (according to the Wuhan YP_009724390.1 sequence), and the spike RBDs as sequenced from the cited VoC, showing an amino acid replacement at the RBD positions K417; N439; L452, E484; S494; N501. The “*” symbols and the labels indicate the SARS-CoV-2 spike RBD positions involved in an amino acid replacement. The sequence of the SARS-CoV-2 chimeric RBD (showing a three amino acid deletion and 22 missense mutations not present in the investigated VoC) from 6vw1.pdb is reported for comparative purposes.

The replacement of the seven investigated residues (N501Y, yellow sticks; E484K/Q, green/pink sticks; N439K, orange sticks; K417N/T, cyan/teal sticks; L452R, dark-blue sticks; S477N, light pink sticks; S494P, hot-pink sticks; Fig. 2) introduces an important perturbation in the local secondary structure of the “boat-shaped” receptor binding motif (RBM) located on the RBD head, consisting of a “bow” portion (residues 456-459 and 468-490, 6m0j.pdb RBD sequence numbering, Fig. 2), a “hull” portion (residues 450-455 and 491-496, 6m0j.pdb RBD sequence numbering, Fig. 2), and a “stern” portion (residues 436-449 and 497-503, 6m0j.pdb RBD sequence numbering, Fig.2). Notably, the RBM highlighted on the head of the spike RBD responsible for the binding interactions with ACE2, due to the formation of H-bonds and/or ionic and aromatic interactions.

**Figure 2.**
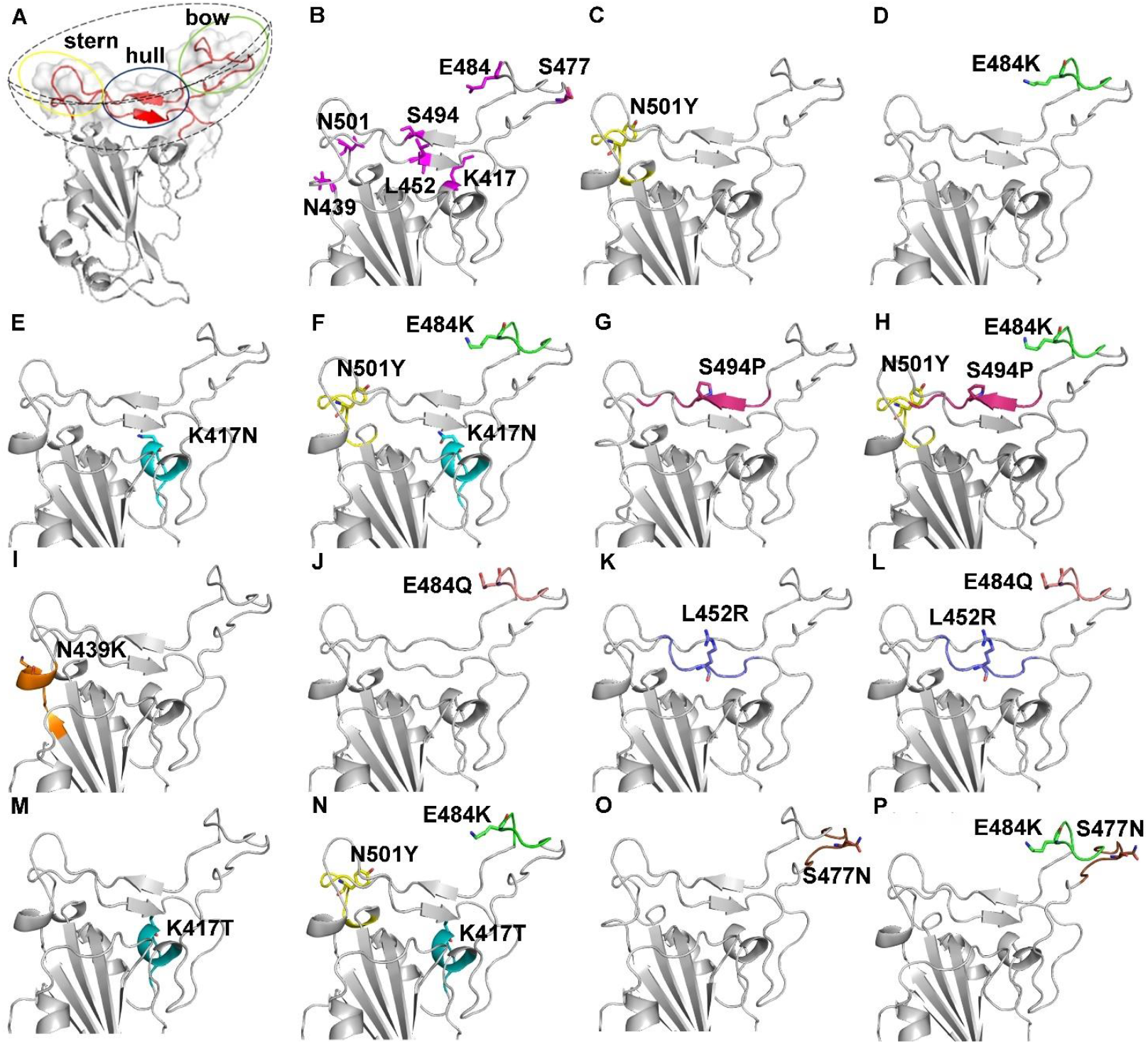
3D comparative models of the investigated RBD mutants. Panle A. The SARS-CoV-2 spike RBD (6m0j.pdb) is reported in white cartoon representation. The RBM is highlighted in red cartoon and transparent surf representation. A boat-shaped dashed line is reported around the RBM for defining the 3 different regions, namely the stern region (yellow circle), consisting of residues 436-449 and 497-503, the hull region (blue circle), consisting of residues 450-455 and 491-496, and the bow region (green circle), consisting of residues 456-459 and 468-490 (6m0j.pdb, RBD sequence numbering). Those three regions delimit the RBM area directly involved in binding interactions with ACE2. All the investigated VoC show missense mutations at the 3 highlighted regions within the RBM. Panel B. Zoomed view of the RBD (white cartoon) showing the RBD amino acids K417; N439; L452; S477; E484; S494; N501 (magenta sticks) observed mutated in the investigated VoC. Panels C-N. Zoomed views of the RBD investigated mutants: N501Y, yellow sticks (C); E484K, green sticks (D); K417N cyan sticks (E), the triple mutant K417N_E484K_N501Y observed in the P.1-Japan/Brazil VoC (F); S494P in dark pink sticks (G); the triple mutant E484K_S494P_N501Y detected in the B.1.1.7_UK VoC (H); N439K, orange sticks, observed in the B.1.141 VoC (I); E484Q, light pink sticks (J); L452R in dark-blue sticks, observed in the B.1.427 California VoC (K); the double mutant E484Q_L452R observed in the B.1.617.1 India VoC (L); K417T in teal sticks (M); the triple mutant N501Y_E484K_K417T, observed in the B.1.351. SA VoC (N); S477N, brown sticks (O); the two RBD mutations S477N_E484K, observed in the B.1.620 VoC (P).

More in detail, the investigated L452R and E484Q amino acid replacements cause a local re-arrangement that perturbs the small beta-sheet in the “hull” region of the boat-shaped RBM (residues 450-455, Fig. 2), whereas N501Y, K417N, E484K and N439K cause a conformational change in an alpha-helix close to N501 and N439 in the “stern” region (residues 436-449 and 497-503). Although S494P, approximately located in the center of the “boat-hull” region (residues 491-496), and S477N, located on the tip of the “boat-bow” region (residues 468-490) do not produce a viewable local conformational change, it is well known that the replacement of a serine may confer a different flexibility to the local secondary structure elements hosting the investigated mutation, due to the different abilities of Ser/Pro/Asn/Thr residues in producing kink/hinge movements [44, 45].

The perturbation of the local secondary structure of structural elements hosting the cited amino acid replacements also triggers the formation of new H-bonds, ionic and aromatic interactions and local conformational changes at the protein-protein interface along interactions with ACE2 (Fig. 3, Fig. 4, Tab. 1, Supp. Tab. 1).

Indeed, it is possible to count a slight increase in the hydrophobic interactions within a range of 5 Å at the SARS-CoV-2 spike RBD/ACE2 interface in the B.1.1.7-UK (S494P_N501Y_E484K); in the B.1.351-SA (N501Y_E484K_K417T) or in the P1. Japan/Brazil (N501Y_E484K_K417N) VoC (Supp. Table 1 and Fig. 4).

**Figure 3.**
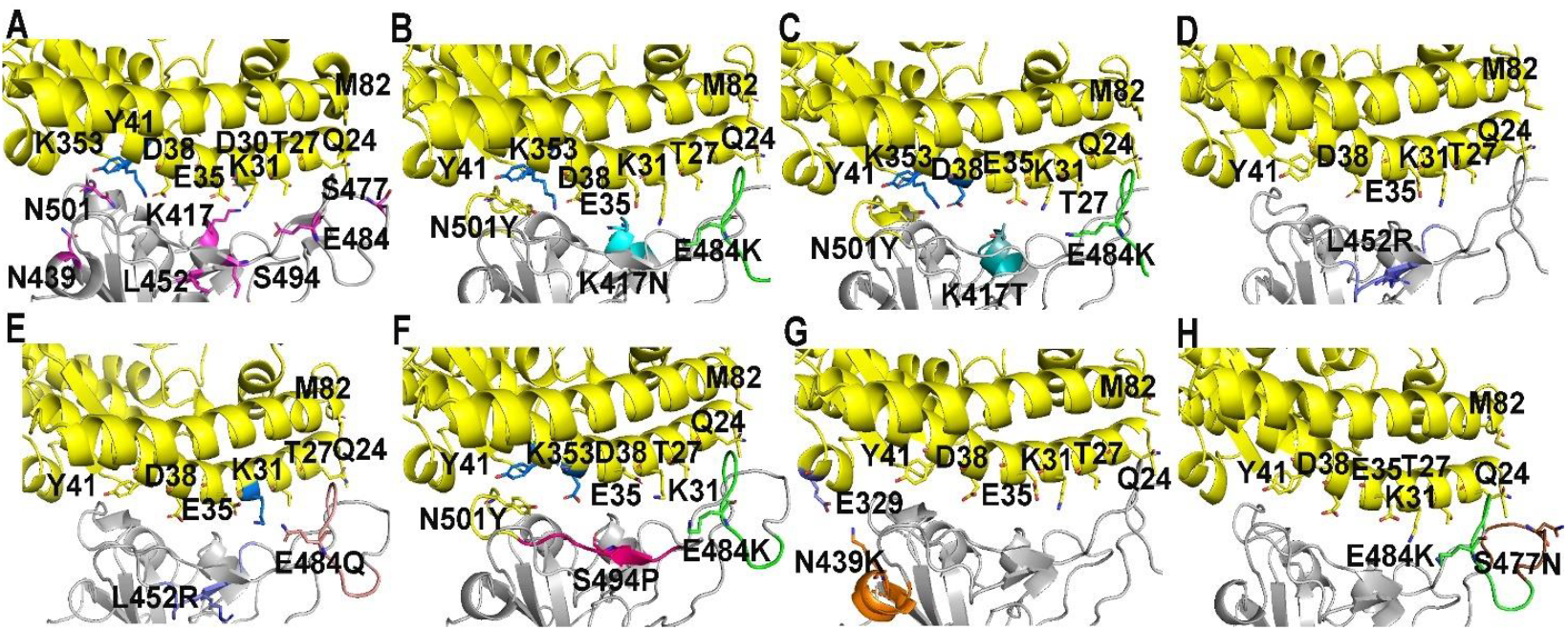
SARS-CoV-2 spike RBD–ACE2 interactions in presence of the investigated amino acid replacements. Panel A. Zoomed view of ACE2 (yellow cartoon) interactions with SARS-CoV-2 spike RBD head (white cartoon) are reported. Amino acid positions involved in the investigated mutants are reported in magenta sticks. Panels B-G. Zommed views of ACE2 interactions with the SARS-CoV-2 spike RBD head of the P.1-Japan/Brazil VoCs (B), B.1.351-S. Africa VoC (C); B.1.427 California VoC (D); B.1.617.1 India VoC (E); B.1.1.7-UK VoC (F); B.1.141 VoC (G); B.1.620 VoC (H). ACE2 is reported in yellow cartoon in all the panels. ACE2 residues within 4 Å from the entire RBD are reported in yellow sticks, whereas ACE2 residues within 4 Å from the investigated RBD mutations are reported in blue sticks in all the panels.

**Figure 4.**
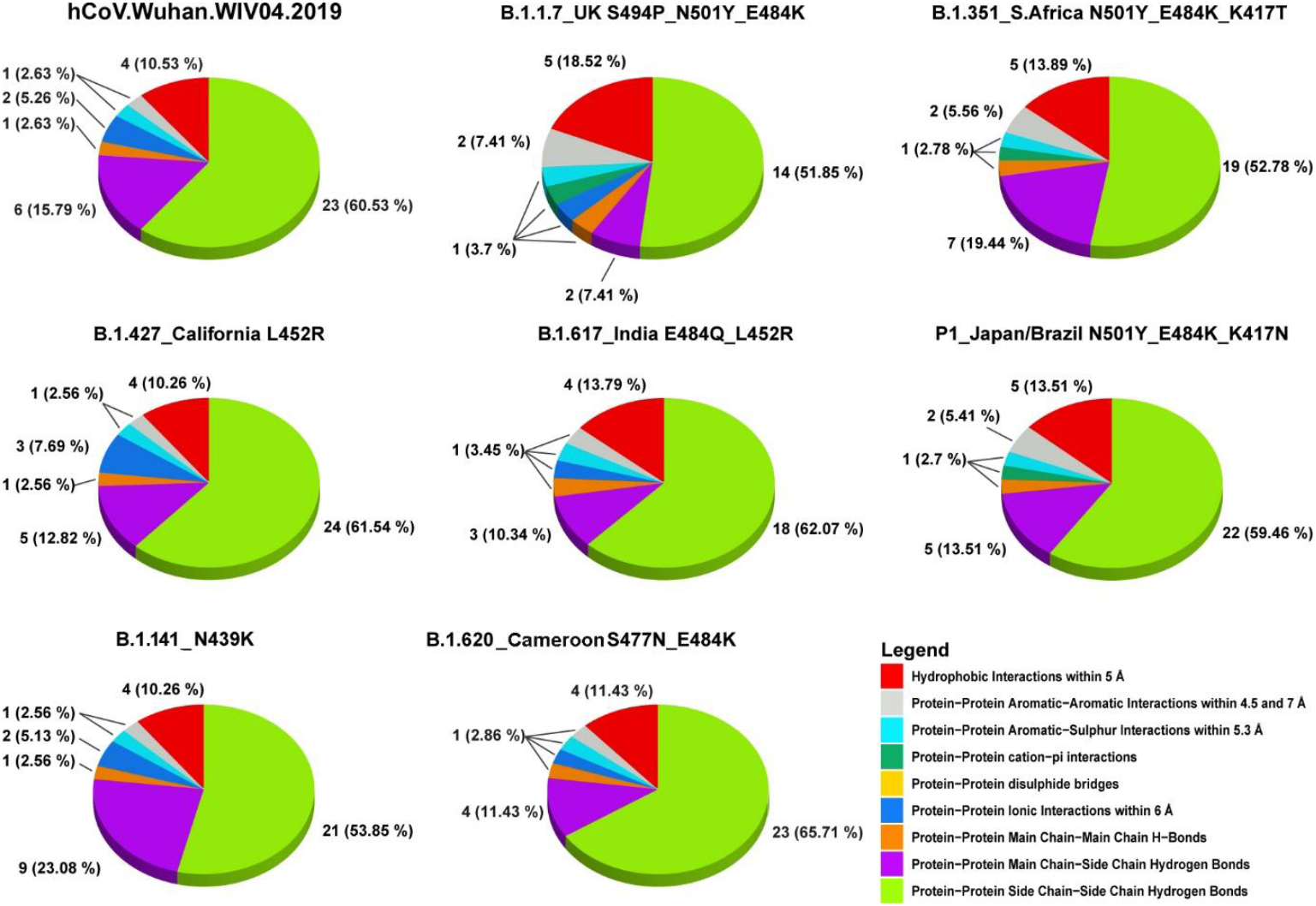
Pie charts summarize the SARS-CoV-2 spike RBD/ACE2 interactions detected by PiC analysis. The pie charts are realized with R (version 4.0.5) by means of ggplot2 library.

Some variations are observed in the number of Protein-Protein Main Chain-Side Chain Hydrogen Bonds which are apparently decreased at the SARS-CoV-2 spike RBD/ACE2 interface in the B.1.1.7-Uk (S494P_N501Y_E484K), in the B1.617-India (E484Q_L452), in the B.1.427 California (L452R), and in the B1.617. Indian (E484Q_L452R) VoC, whereas the same interactions appear increased in number in the B.1.351-SA (N501Y_E484K_K417T) and in the B1.141 VoC N439K (Supp. Tab. 1 and Fig. 4).

A larger number of variations are observed in the number of Protein-Protein Side Chain-Side Chain Hydrogen Bonds. Indeed, side chain – side chain H-bonds decrease at the SARS-CoV-2 spike RBD/ACE2 interface in all the VoC with the exception of the B.1.427 California L452R VoC (Supp. Tab. 1 and Fig. 4).

New Protein-Protein Aromatic-Aromatic Interactions (within 4.5 and 7 Å) and Protein-protein cation-pi interactions are observed at the SARS-CoV-2 spike RBD/ACE2 interface in all the VoCs showing the N501Y amino acid replacement (with specific reference to B.1.1.7-Uk (S494P_N501Y_E484K), B.1.351-SA (N501Y_E484K_K417T) and P1. Japan/Brazil (N501Y_E484K_K417N) showing new aromatic interactions with the ACE2 Tyr41 and cation-pi interactions with the ACE2 Lys353 (Supp. Tab. 1 and Fig. 4). The number of Protein-Protein Ionic Interactions (within 6 Å) at the SARS-COV-2 spike RBD/ACE2 interface may increase depending on the investigated amino acid replacement and its steric hinderance. I.e., L452R is the mutant with the greater number of Protein-Protein ionic interactions (Supp. Tab. 1 and Fig. 4).

From an energetical point of view the B.1.351-SA VoC, showing the three mutations N501Y_E484K_K417T at the RBD, has the highest binding affinity (−21,37 Kcal/mol, Table 1) for ACE2 (increased of 4% with respect to the Wuhan spike RBD, -20.51 Kcal/mol, Table 1), followed by the B.1.141-VoC showing the single N439K amino acid replacement at the RBD and the single mutant E484K firstly detected in the spike RBD of the B.1.351-SA VoC (Table 1). All the other single and multiple amino acid replacements show a slightly decreased interaction energy with ACE2 (Table 1), at variance with what observed for the Wuhan SARS-CoV-2 spike RBD/ACE2 interactions.

**Table 1.** FoldX interection energies between the ACE2 human receptor and the investigated RBD amino acid replacements,

|  | Wuhan SARS-CoV-2 Spike/ ACE2 (6m0j.p db) | N501Y | E484K | K417N | K417T | S494P | E484Q | S477N | B.1.427 California (L452R) | B1.141 N439K | P.1. Japan/ Brazil (N501Y_ E484K_ K417N) | B.1.617. 1 India (E484Q_ L452R) | B.1.351. SA (N501Y_ E484K_ K417T) | B.1.1.7 UK (S494P_ N501Y_ E484K) | B.1.620 (S477N_ E484K) |
| --- | --- | --- | --- | --- | --- | --- | --- | --- | --- | --- | --- | --- | --- | --- | --- |
| Group1 (ACE2) | A | A | A | A | A | A | A | A | A | A | A | A | A | A | A |
| Group2 (RBD) | E | E | E | E | E | E | E | E | E | E | E | E | E | E | E |
| Interaction Energy (Kcal/mol) | **<u>-20,51</u>** | -19,60 | **<u>-20,67</u>** | -19,38 | -18,73 | -19,37 | -19,03 | -18,96 | -20,16 | **<u>-20,78</u>** | -18,80 | -18,86 | **<u>-21,37</u>** | -18,12 | -18,26 |
| IntraclashesGroup1 | 22,29 | 21,70 | 25,14 | 21,20 | 23,84 | 22,98 | 23,69 | 25,03 | 24,12 | 20,69 | 23,13 | 24,44 | 21,94 | 27,75 | 24,71 |
| IntraclashesGroup2 | 5,49 | 4,34 | 7,10 | 7,53 | 5,47 | 5,51 | 8,10 | 5,58 | 6,60 | 6,35 | 7,27 | 5,73 | 6,52 | 5,81 | 5,57 |
| Backbone Hbond | -3,64 | -3,01 | -3,90 | -2,58 | -2,71 | -2,89 | -2,84 | -2,80 | -2,67 | -2,67 | -3,08 | -3,27 | -2,95 | -1,72 | -2,81 |
| Sidechain Hbond | -12,12 | -12,85 | -12,89 | -10,55 | -9,39 | -7,09 | -8,93 | -11,03 | -10,77 | -12,03 | -11,57 | -8,52 | -11,59 | -7,09 | -10,67 |
| Van der Waals | -15,89 | -16,63 | -16,28 | -15,44 | -15,59 | -15,51 | -15,70 | -15,74 | -15,92 | -15,78 | -16,54 | -15,13 | -15,84 | -15,64 | -16,13 |
| Electrostatics | -2,41 | -2,79 | -2,65 | -1,82 | -1,90 | -2,55 | -2,42 | -2,28 | -3,05 | -2,50 | -1,10 | -2,32 | -1,11 | -2,21 | -2,49 |
| Solvation Polar | 22,14 | 23,39 | 22,95 | 20,34 | 20,79 | 21,07 | 20,61 | 21,90 | 22,65 | 21,56 | 21,33 | 19,78 | 20,33 | 19,91 | 22,58 |
| Solvation Hydrophobic | -20,01 | -20,93 | -20,30 | -19,62 | -19,81 | -19,90 | -19,88 | -19,80 | -19,85 | -20,18 | -21,33 | -19,45 | -20,38 | -20,56 | -20,13 |
| Van der Waals clashes | 0,41 | 0,81 | 0,33 | 0,36 | 0,23 | 0,38 | 0,57 | 0,35 | 0,42 | 0,42 | 0,43 | 0,39 | 0,45 | 0,78 | 0,32 |
| entropy sidechain | 9,53 | 10,72 | 10,13 | 8,75 | 8,84 | 6,25 | 8,54 | 9,78 | 7,59 | 9,81 | 10,56 | 7,87 | 9,05 | 7,43 | 9,66 |
| entropy mainchain | 1,57 | 1,78 | 1,70 | 1,09 | 0,77 | 1,04 | 1,24 | 0,80 | 1,73 | 0,85 | 2,34 | 1,67 | 0,70 | 1,15 | 1,68 |
| torsional clash | 0,08 | 0,06 | 0,42 | 0,08 | 0,08 | 0,07 | 0,08 | 0,09 | 0,06 | 0,05 | 0,17 | 0,37 | 0,05 | 0,10 | 0,07 |
| backbone clash | 2,49 | 2,20 | 2,57 | 1,84 | 2,23 | 2,20 | 2,23 | 2,25 | 2,14 | 2,49 | 2,33 | 2,18 | 2,30 | 2,00 | 2,24 |
| helix dipole | -0,06 | -0,04 | 0,01 | -0,10 | -0,11 | -0,06 | -0,09 | -0,12 | -0,10 | -0,02 | -0,04 | -0,09 | -0,12 | -0,03 | -0,12 |
| disulfide | 1,78E-15 | 3,55E-15 | -1,78E-15 | -7,11E-15 | 0,00E+00 | -3,55E-15 | 3,55E-15 | 1,78E-15 | -7,11E-15 | -1,78E-15 | -3,55E-15 | 0,00E+00 | -1,78E-15 | 5,33E-15 | -5,33E-15 |
| electrostatic kon | -0,18 | -0,18 | -0,26 | 0,05 | -0,01 | -0,17 | -0,27 | -0,16 | -0,32 | -0,36 | -0,04 | -0,20 | -0,03 | -0,32 | -0,29 |
| energy Ionisation | 0,06 | 0,06 | 0,07 | 0,06 | 0,05 | 0,00 | 0,05 | 0,06 | 0,05 | 0,06 | 0,06 | 0,05 | 0,05 | 0,06 | 0,06 |
| Entropy Complex | 2,38 | 2,38 | 2,38 | 2,38 | 2,38 | 2,38 | 2,38 | 2,38 | 2,38 | 2,38 | 2,38 | 2,38 | 2,38 | 2,38 | 2,38 |
| Number of Residues | 791 | 791 | 791 | 791 | 791 | 791 | 791 | 791 | 791 | 791 | 791 | 791 | 791 | 791 | 791 |

With the exception of the B.1.427 California VoC (L452R) and B.1.141 (N439K) showing one more interaction (39 interactions) with respect to the interactions detected at the Wuhan SARS-CoV-2 spike RBD/ACE2 protein-protein interface (38 residues), all the other investigated ACE2/RBD complexes showed a decrease in the number of detected interactions at the protein-protein interface caused by the investigated amino acid replacements. The B.1.1.7-UK (S494P_N501Y_E484K) VoC and the B1.617-India (E484Q_L452R) VoC show the lowest number of interactions (27 and 29, respectively) at the RBD/ACE2 protein-protein interface, according to PIC estimations (Fig. 4 and Supp. Tab. 1).

### Sampling of ACE2 structurally related alternative receptors via folding recognition and multiple sequence alignments (MSA)

ACE2 sequence was used for screening the PDB, searching for ACE2 structurally related proteins that might work as host-cell entry sites for SARS-CoV-2. The performed screening revealed the presence of several crystallized structures of ACE2 and of the structural/functional related ACE protein from several mammalia and insecta species (Table 2). Furthermore, the screening revealed the presence of several oligopeptidases from bacteria and protista, structurally related to ACE2 in the PDB (Supp. Tab. 2). Notably, two mammalia oligopeptidases, THOP1 and NLN, were highlighted as ACE2 structurally related proteins (Tab. 2). ACE2 [46] shares with ACE [47], THOP1 [48], and NLN [49] the 34% (49%), 18% (33%) and 17% (35%) of identical (similar) residues.

**Table 2.**
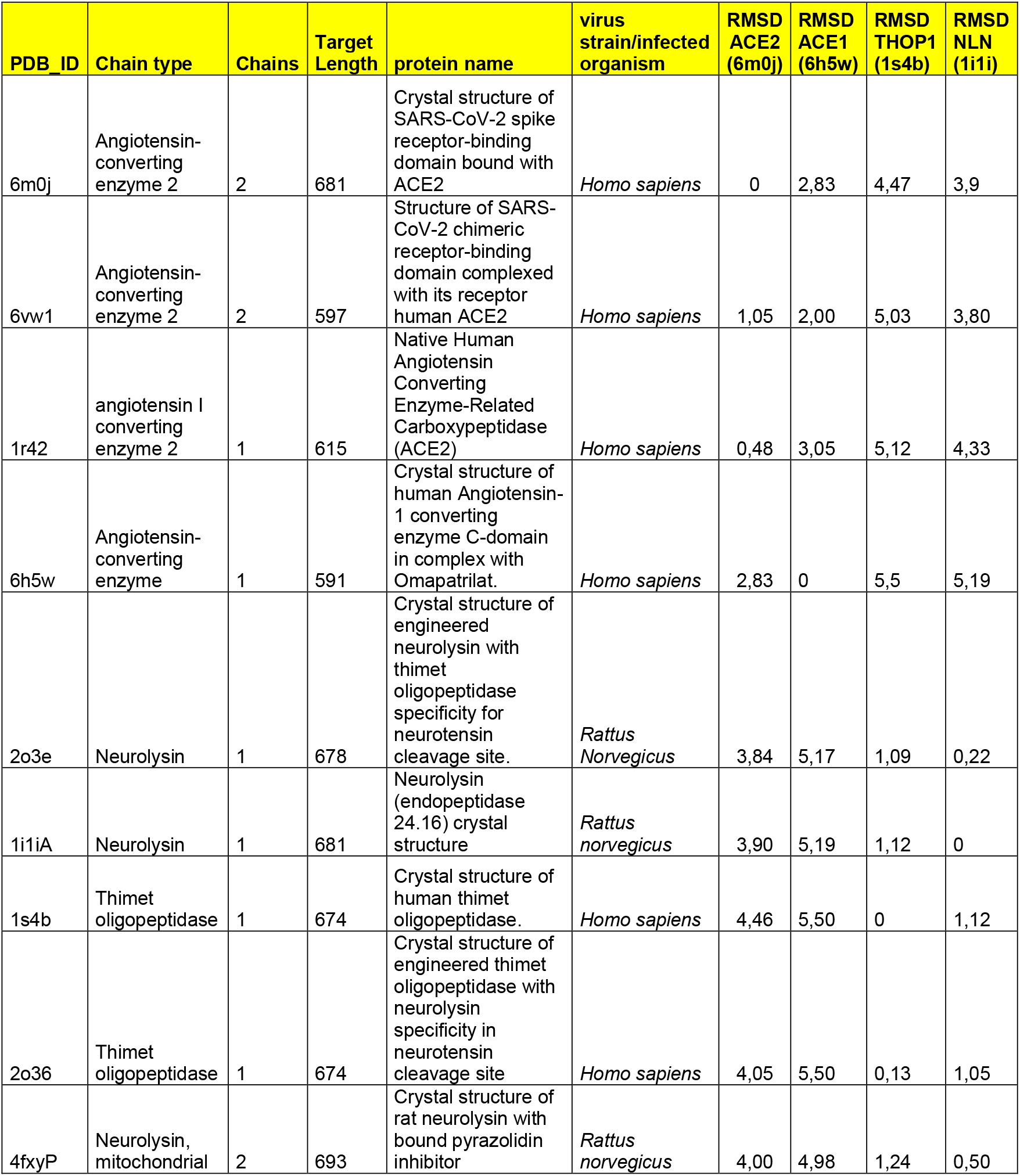
Extract of ACE2 structurally/functionally related proteins sampled by pGenThreader/ITASSER. (see also Supp. Tab. 2)

| PDB_ID | Chain type | Chains | Target Length | protein name | virus strain/infected organism | RMSD ACE2 (6m0j) | RMSD ACE1 (6h5w) | RMSD THOP1 (1s4b) | RMSD NLN (1i1i) |
| --- | --- | --- | --- | --- | --- | --- | --- | --- | --- |
| 6m0j | Angiotensin-converting enzyme 2 | 2 | 681 | Crystal structure of SARS-CoV-2 spike receptor-binding domain bound with ACE2 | *Homo sapiens* | 0 | 2,83 | 4,47 | 3,9 |
| 6vw1 | Angiotensin-converting enzyme 2 | 2 | 597 | Structure of SARS-CoV-2 chimeric receptor-binding domain complexed with its receptor human ACE2 | *Homo sapiens* | 1,05 | 2,00 | 5,03 | 3,80 |
| 1r42 | angiotensin I converting enzyme 2 | 1 | 615 | Native Human Angiotensin Converting Enzyme-Related Carboxypeptidase (ACE2) | *Homo sapiens* | 0,48 | 3,05 | 5,12 | 4,33 |
| 6h5w | Angiotensin-converting enzyme | 1 | 591 | Crystal structure of human Angiotensin-1 converting enzyme C-domain in complex with Omapatrilat. | *Homo sapiens* | 2,83 | 0 | 5,5 | 5,19 |
| 2o3e | Neurolysin | 1 | 678 | Crystal structure of engineered neurolysin with thimet oligopeptidase specificity for neurotensin cleavage site. | *Rattus Norvegicus* | 3,84 | 5,17 | 1,09 | 0,22 |
| 1i1iA | Neurolysin | 1 | 681 | Neurolysin (endopeptidase 24.16) crystal structure | *Rattus norvegicus* | 3,90 | 5,19 | 1,12 | 0 |
| 1s4b | Thimet oligopeptidase | 1 | 674 | Crystal structure of human thimet oligopeptidase. | *Homo sapiens* | 4,46 | 5,50 | 0 | 1,12 |
| 2o36 | Thimet oligopeptidase | 1 | 674 | Crystal structure of engineered thimet oligopeptidase with neurolysin specificity in neurotensin cleavage site | *Homo sapiens* | 4,05 | 5,50 | 0,13 | 1,05 |
| 4fxyP | Neurolysin, mitochondrial | 2 | 693 | Crystal structure of rat neurolysin with bound pyrazolidin inhibitor | *Rattus norvegicus* | 4,00 | 4,98 | 1,24 | 0,50 |

The four proteins share a very similar overall structure (Fig. 5). Indeed, the RMSD of the atomic coordinates of the available THOP1 and NLN crystallized structures and the investigated ACE2 structure ranges between 3.9 and 4.5 Å (Tab. 2, Fig. 5).

**Figure 5.**
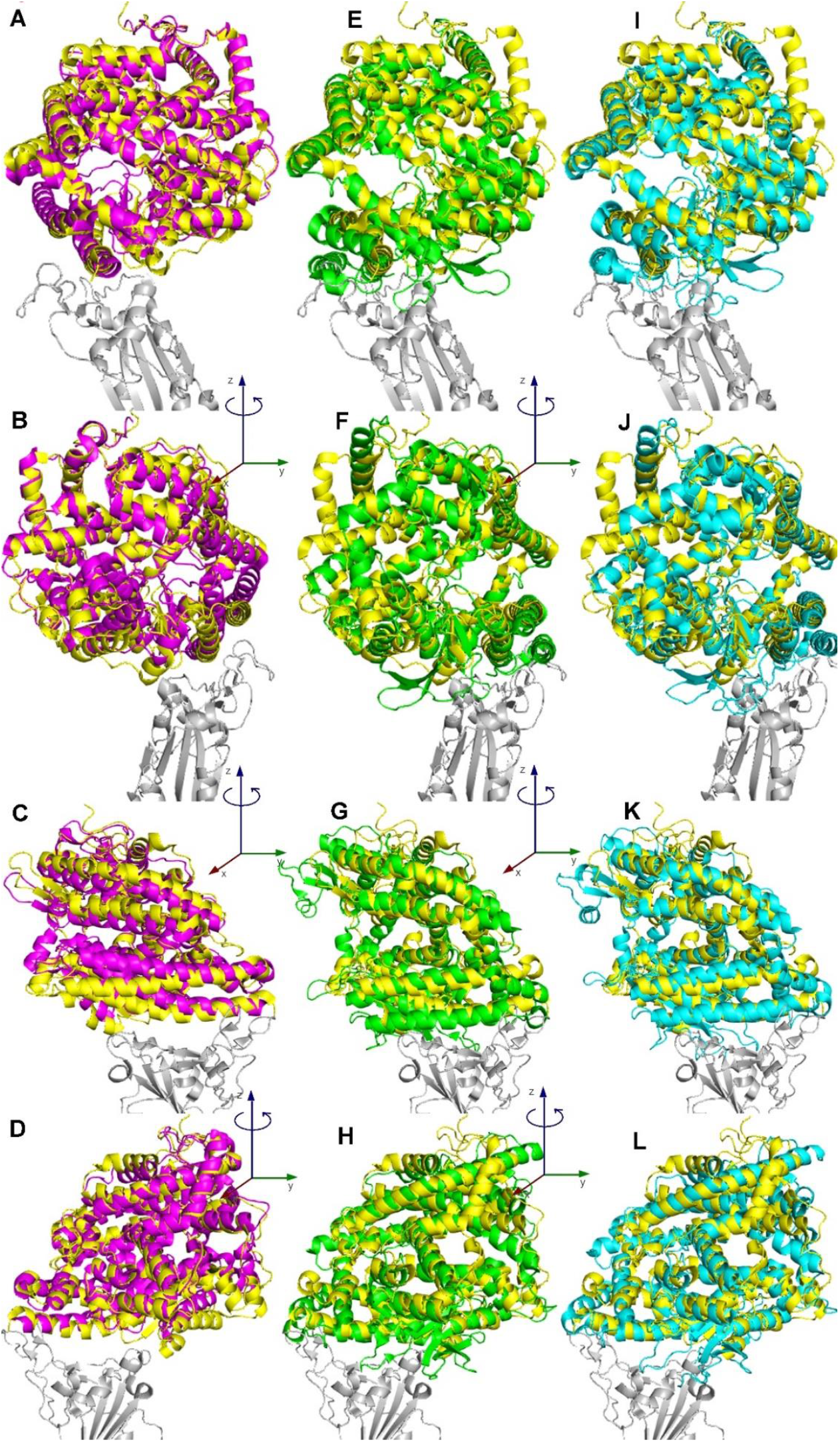
Superimposition of the ACE2 structurally related proteins ACE, THOP1 and NLN to ACE2. View of the lateral view rotated of 90 degrees (along the RBD z axis) along each column showing the superimposition of ACE (6h5w.pdb, magenta cartoon, Panels A-D), THOP1 (1s4b.pdb, green cartoon, Panels E-H) and NLN (1i1i.pdb, cyan cartoon, Panels I-L) on the structure of the ACE2 receptor (6m0j.pdb, yellow cartoon) interacting with the SARS-CoV-2 spike RBD (6m0j.pdb, light grey cartoon).

### SARS-CoV-2 spike RBD variant interactions with the ACE2 structurally related proteins: a comparative analysis (number of interactions, local secondary structure perturbation, and interaction energy calculation)

At the protein-protein interface it is possible to see that ACE shows two alpha-helices, similarly oriented to the two alpha-helices that represent in ACE2 the main surface of interaction with SARS-CoV-2 spike RBD. THOP1 and NLN show the same alpha-helices in close contact with the SARS-CoV-2 spike RBD and an extra helix parallel to the previous two, forming other interactions with the RBD of the SARS-CoV-2 spike RBD (Fig. 6).

**Figure 6.**
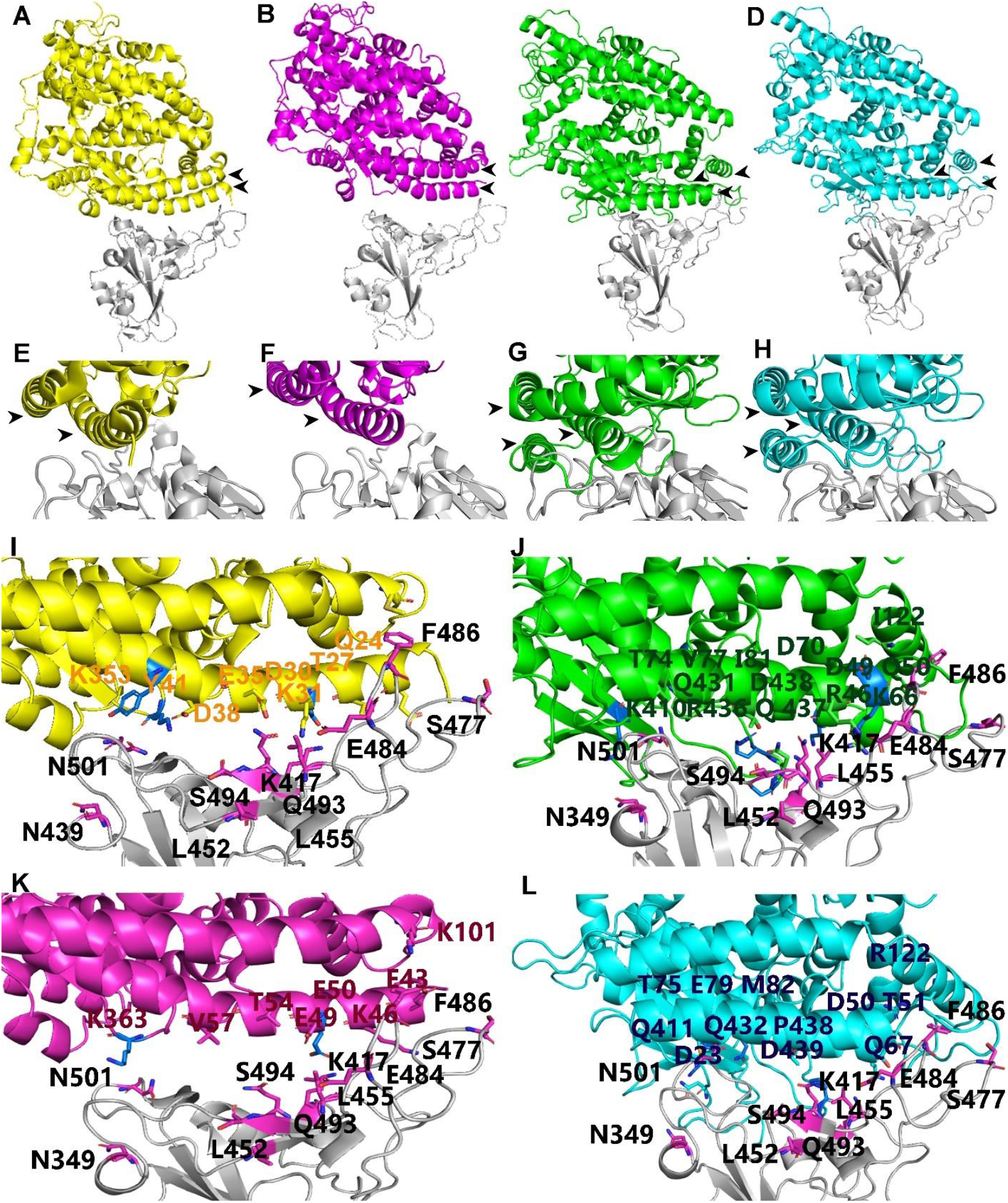
Comparative structural analysis of the ACE2 structurally related proteins ACE, THOP1 and NLN. The SARS-CoV-2 spike RBD (light grey cartoon) complexed with ACE2 (panel A; yellow cartoon), ACE (panel B, magenta cartoon), THOP1 (panel C, green cartoon) and NLN (panel D, cyan cartoon). Panels E-H. Zoomed views of the SARS-CoV-2 spike RBD interacting with ACE2 (E), ACE (F), THOP1 (G), and NLN (H). Black arrow-heads in panels A-H indicate secondary structure elements mainly involved in interactions with the SARS-CoV-2 spike RBD. Panels I-L. Zoomed view of the RBD (light grey cartoon) showing the RBD amino acids K417; N439; L452; E484; S494; N501 (magenta sticks) observed mutated in the investigated VoC. Residues within 4 Å from SARS-CoV-2 spike RBD highlighted within ACE2 (I, yellow), ACE (K, magenta), THOP1 (J, green), and NLN (L, cyan) are reported in coloured sticks. ACE2 structurally related receptor residues within 4 Å from the protein positions observed mutated in the investigated VoCs are reported in bluse sticks (see Supp. Tab. 1 for a list of the detected interactions between the Wuhan SARS-CoV-2 spike RBD and ACE2 structurally related receptors according to PIC estimations).

Although the investigated proteins share a low percentage of identical residues, local secondary structures crucial for interactions with the SARS-CoV-2 spike RBD, consisting of the investigated receptor residues located within 4 Å from the RBD are very similar (Fig. 7). The ACE protein region in contact with the RBD is the most similar to the corresponding counterpart in ACE2 (Fig. 7). The four proteins share one longer helix similarly oriented (residues L29-Q50, ACE2 sequence numbering; and residues L65-Q86, THOP1 sequence numbering) and one superimposable beta-sheet different in length (residues T347-L359, ACE2 sequence numbering, and residues A426-L459, THOP1 sequence numbering). THOP1 and NLN show an extra helix consisting of their N-terminal region (residues L24-T64, THOP1 sequence numbering) and show a beta-sheet in place of the helix located close to the “stern” portion of the boat-shaped RBD head (residues T324-L333, ACE2 sequence numbering, and residues E406-L415, THOP1 sequence numbering). A last region with a relative different orientation in the space consists of the THOP1 bent helix (residues M112-K128, THOP1 sequence numbering) corresponding to the ACE2 bent helix (residues L73-Q89, ACE2 sequence numbering) (Fig. 7).

**Figure 7.**
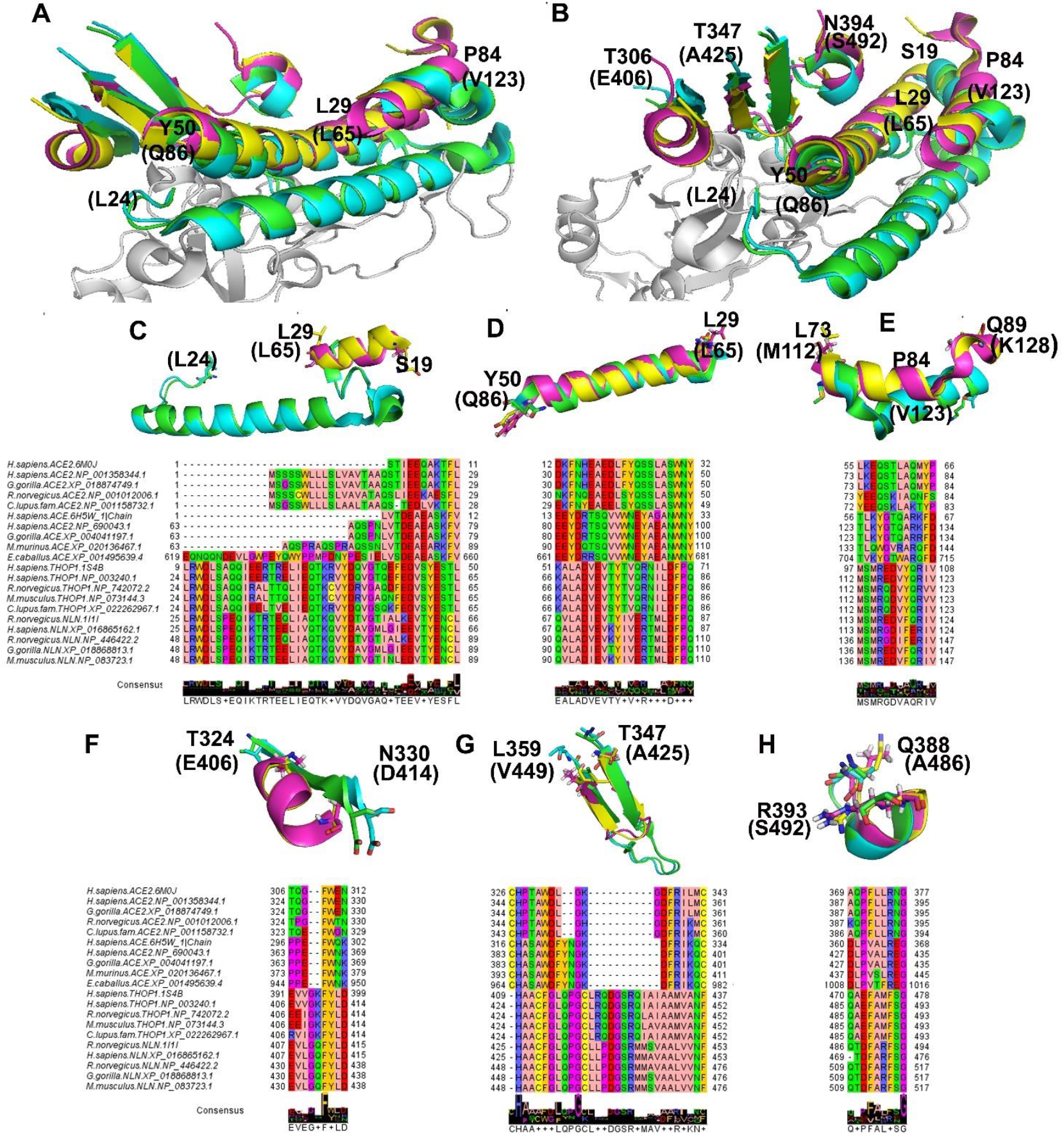
Zommed views of the SARS-CoV-2 spike RBD head in contact with the ACE2 structurally related proteins. Panels A-B. The RBD head portion is reported in light grey cartoon. ACE2, ACE, THOP1 and NLN are reported in yellow, magenta, green and cyan cartoon, respectively. Panels C-H. Zoomed view of secondary structural elements highlighted from ACE2 structurally related proteins. Residues ad the beginning and at the end of the reported secondary structural elements are indicated by labels. Multiple sequence alignment of the highlighted secondary structural elements are reported below each highlighted structural element of panels C-H.

While ACE shows an interaction energy with RBD lower than the ones observed in presence of ACE2, the calculated interaction energies between the SARS-CoV-spike RBD and THOP1 or NLN appear to be stronger than those calculated for ACE2 (Table 3).

**Table 3.** FoldX interection energies between SARS-CoV-2 RBD and ACE2 structurally related proteins.

| PDB | ACE2<br>(6m0j.pdb) | ACE<br>(6h5w.pdb) | NLN<br>(1i1i.pdb) | THOP1<br>(1s4b.pdb) |
| --- | --- | --- | --- | --- |
| Group1 (Chain) | A (ACE2) | A (ACE1) | P (NLN) | P (THOP) |
| Group2 (Chain) | E (RBD_6m0j) | E (RBD_6m0j) | E (RBD_6m0j) | E (RBD_6m0j) |
| Interaction Energy | **-20,52** | **-7,41** | **-23,72** | **-22,00** |
| IntraclashesGroup1 | 22,28 | 22,95 | 21,50 | 22,95 |
| IntraclashesGroup2 | 5,49 | 6,63 | 13,07 | 9,70 |
| Backbone Hbond | -3,64 | -0,16 | -13,08 | -7,86 |
| Sidechain Hbond | -12,12 | -9,76 | -19,3 | -9,39 |
| Van der Waals | -15,89 | -7,81 | -55,11 | -48,08 |
| Electrostatics | -2,41 | -2,65 | -1,62 | -3,31 |
| Solvation Polar | 22,14 | 12,62 | 84,22 | 76,28 |
| Solvation Hydrophobic | -20,02 | -9,90 | -68,74 | -60,04 |
| Van der Waals clashes | 0,41 | 0,06 | 6,96 | 2,91 |
| entropy sidechain | 9,53 | 9,02 | 18,80 | 18,02 |
| entropy mainchain | 1,57 | 1,33 | 21,16 | 8,86 |
| torsional clash | 0,083 | 0,055 | 3,68 | 0,94 |
| backbone clash | 2,45 | 0,79 | 17,26 | 14,11 |
| helix dipole | -0,06 | -0,02 | -0,00 | 0,14 |
| Disulfide | 1,78E-15 | 3,55E-15 | 0 | 0 |
| electrostatic kon | -0,18 | -0,20 | -0,67 | -0,47 |
| energy Ionisation | 0,07 | -2,84E-16 | 7,06E-03 | 1,60E-16 |
| Entropy Complex | 2,38 | 2,38 | 2,38 | 2,38 |
| Number of Residues | 791 | 780 | 859 | 848 |

### Expression of ACE2 structurally related receptors as coronavirus entry factors

GTEx database was screened for estimating the expression levels of ACE2, ACE, THOP1 and NLN together with other SARS-CoV-2 host-cell entry factors in all the tissues available on GTEx. From this analysis, THOP1 resulted more expressed than ACE2 in all the screened tissues (with the exception of kidney cortex, heart left ventricle, adipose visceral, small intestine, showing THOP1/ACE2 similar expression levels) and, most importantly, it is highly expressed in the lung, in the colon, in the esophagus mucosa and in all the brain compartments, in which ACE2 appears poorly expressed (Fig. 8). The expression of NLN or ACE is comparable to or slightly higher than ACE2 expression in all the investigated tissues with the exception of heart, adipose tissue, intestine and testis.

**Figure 8.**
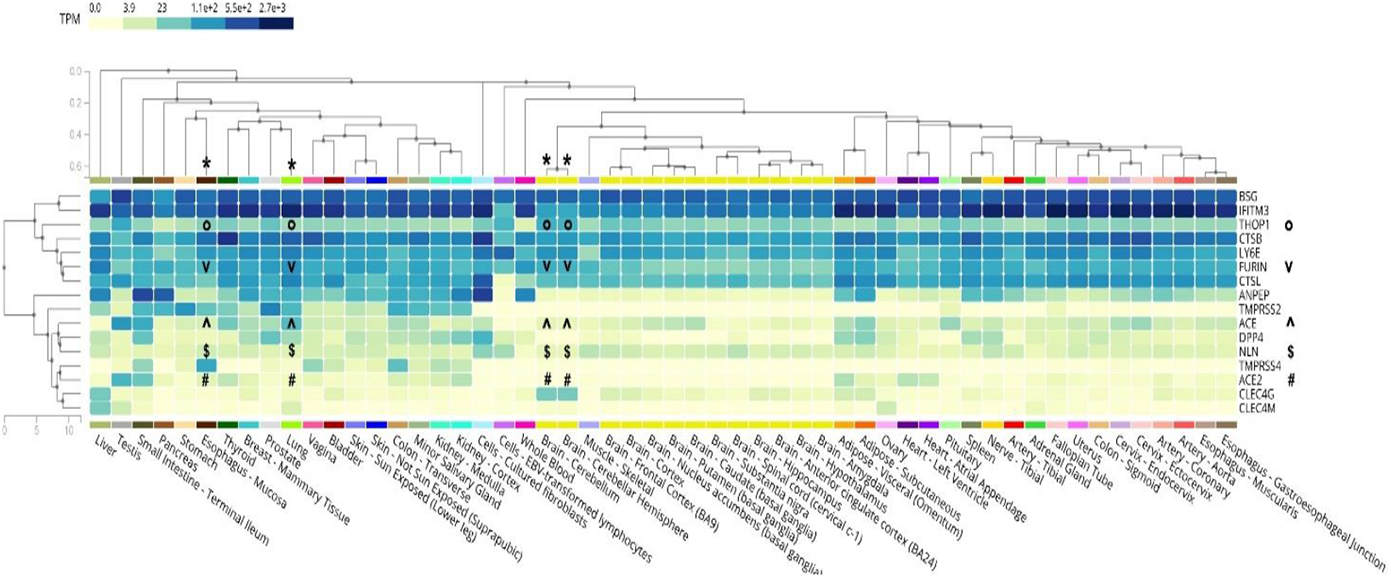
Expression of SARS-CoV-2 entry factors. Heatmap analysis about the expression of several SARS-CoV-2 entry factors and ACE2 structurally related proteins was obtained through GTEx multi-gene query expression (GTEx based on V8 Release). The “*” symbols indicate the position of the main discussed tissues (brain, esophagus mucosa, lung). The “°, #, v, ^, $” symbols indicate THOP1, ACE2, Furin, ACE, NLN expression levels, respectively.

## DISCUSSION

The rapid spread of new SARS-CoV-2 variants [16, 17, 50] makes necessary the development of new tools for evaluating the interactions of SARS-CoV-2 proteins for the host cell receptors. A crucial role in SARS-CoV-2 infection is played by the SARS-CoV-2 spike protein, whose interactions with ACE2 receptor triggers pre-/post-fusion conformational changes causing the virus entry into the human cells [1, 9]. The spike domain responsible for direct interactions with the human ACE2 receptor is the RBD [1, 3, 41, 51–53]. Thus, great attention is dedicated to mutations occurring at the SARS-CoV-2 spike RBD, because they can cause an increase in the binding affinity of the spike protein for the human ACE2 receptor, which may reflect an increased transmissibility or a new acquired antibody escape ability [1, 12–15, 54].

Previously [1], we have determined the interaction energies between the SARS-CoV-2 spike RBD and the human ACE2 receptor available under the PDB_ID 6vw1.pdb [1, 10]. However, the crystallized structure of 6vw1.pdb consisted of a chimeric SARS-CoV-2 spike RBD interacting with ACE2 [10]. The chimeric RBD from 6vw1.pdb showed 22 missense mutations and the deletion of three residues at the RBD with respect to the amino acid sequence of SARS-CoV-2 spike RBD according to the Wuhan reference sequence available under the refseq accession number YP_009724390.1. Less than one year ago coordinates of the SARS-CoV-2 spike RBD (according to YP_009724390.1) crystallized in complex with the human ACE2 receptor were made available under the PDB_ID 6moj.pdb [11]. Thus, we used the 3D coordinates of 6m0j for re-estimating the interaction energies at the SARS-CoV-2 spike RBD/ACE2 protein-protein interface and we modelled RBD mutants on the 6m0j.pdb protein template and quantified the affinity of the known RBD mutants for the human ACE2 receptor. More in detail we studied the impact of the mutations responsible for the investigated VoC B.1.1.7-UK (N501Y, S494P, E484K), P.1-Japan/Brazil (K417N/T, E484K, N501Y), B.1.351-SA (K417N, E484K, N501Y), B.1.427/B.1.429-California (L452R), the B1.141 (N439K) and the recent B1. 617-India (L452R, E484Q) VoC. The effect of the investigated mutations on the SARS-CoV-2 spike RBD structure and on RBD/ACE2 interactions was checked revealing that all of them might perturb the RBD structure either at the level of the secondary structure elements hosting the investigated mutations or at the level of spatially close secondary structure elements on the entire RBD head.

In terms of binding interactions it is possible to see that H-bonds and hydrophobic interactions among side-chains at the protein-protein interface, but also backbone-side chain H-bonds, among the very short-range (<3.8 Å) interactions [55], can substantially increase or decrease as a consequence of a mutation, as observed in the single N439K amino acid replacement of the B1.141 VoC or in the B.1.351-SA VoC consisting of the triple mutant N501Y_E484K_K417T, with respect to the interactions calculated in the crystallized SARS-CoV-2 spike RBD/ACE2 protein complex used as a reference structure. The replacement of N501 with a tyrosine introduces new aromatic-aromatic and π-aromatic binding interactions in the short/medium-range (3.8-9.5 Å) interactions [55].

In terms of binding affinity and interaction energies, it appears that the single amino acid replacements N439K (B1.141 VoC) and E484K (detected in several VoC) cause the most dramatic increase in interaction energies. In addition, the triple mutant N501Y_E484K_K417T detected in the B.1.351-SA VoC [14, 21] shows an increase of about 4 % in binding affinity that might reflect the greater ability shown by this VoC in escaping antibodies produced as a consequence of SARS-CoV-2 Wuhan-sequence based vaccination [14]. Also, the B.1.1.7_UK S494P_N501Y_E484K VoC shows an important variation in the interactions at the ACE2/RBD interface and a decrease in the calculated interaction energies. Notably, this VoC shows the replacement of S494 with a proline residue. P, S, and G play a hinge role in local secondary structures especially when other P, S or G are near in the sequence [45, 56]. The replacement of a serine with a proline may confer less flexibility to the local secondary structure and thus it is retained this mutation limits RBD local flexibility causing a decrease in the number of short/medium-range interactions [55] at the ACE2/RBD interface. It can be argued that the limited flexibility introduced by the S494P amino acid replacement decreases the affinity for ACE2 and makes sensitive this VoC to antibodies produced by vaccination [21], despite of the presence of N501Y and E484K that as single mutants show a higher affinity for the ACE2 receptor. A similar reduced flexibility might be expected for the RBD of the double mutant S477N_E484K due to the replacement of the Ser 477 with an Asn residue.

While it is expected that mutations at the RBD need to be monitored because those mutations can be responsible for an increase in the binding affinity for the ACE2 receptor, it is matter of debate the increased efficiency in entering host-cells proposed for variants showing supplementary mutations far from the RBD, i.e., D614G or P681H shown by several VoC [57] and other variants of interest or under monitoring (https://www.ecdc.europa.eu/en/covid-19/variants-concern). It is retained that mutations like D614G and P681H, being located in the N-terminal portion of the spike pre-fusion conformation, despite of their relatively great distance from the SARS-CoV-2 spike RBD/ACE2 protein-protein interface, can confer a different flexibility (increased for the D614G mutant and decreased for the P681H mutant) to the entire spike protein, before cleavage events determining the post-fusion conformation, following the spike N-terminal loss [1]. Indeed, it was observed that a successful interaction of the spike protein with the ACE2 receptor is supported by the high flexibility of the SARS-CoV-2 spike protein in the pre-fusion conformation, while scanning host-cell surface [2, 3].

Our conclusions about changes in binding energies and number of interactions for the investigated VoC are coherent with the expected effects about spike RBD/ACE2 interactions and antibody escape reported for the described VoC [12–15, 18–21].

While several studies about *in vitro* ACE2/spike binding assays are becoming available, an *in vitro* golden standard technique for quickly estimating the interaction energies between ACE2 and RBD variants is difficult to establish due to relative differences observed between the binding affinity estimated for the ACE2/Wuhan spike protein-protein complex and those estimated for ACE2/RBD variants ([58–60] and Supp. Tab. 3). The observed differences can be ascribed to the employment of the only RBD instead of the entire spike protein, as well as to the employment of living cells expressing ACE2 (full length) or recombinant truncated human ACE2 ectodomain. Also, the different employed ACE2/RBD ratio in the performed binding assays, as well as the different probes and employed detectors can be related to the observed differences in binding affinity (Supp. Tab. 3).

Our *in silico* molecular framework allows to obtain a quick estimation of ACE2/RBD interaction energies that can provide a reliable idea of the ACE2/RBD variant affinity. The same molecular framework can be easily adapted, also based on the future available solved structures, for estimating ACE2/spike interactions by using the entire spike protein instead of the only RBD, for using various ACE2/spike ratios (i.e., 2 spike proteins with 6 ACE2 proteins, as previously described [1]) and to relax the obtained all-atom systems by using molecular dynamics [61, 62].

Among the investigated VoC-RBDs, the triple RBD mutant of the B.1.315 SA VoC shows an increase of less than 5% in the binding affinity compared to the binding affinity estimated for the Wuhan spike RBD/ACE2 protein complex. All the other mutants show lower increases or decreases in the calculated interaction energies, compared to the Wuhan spike RBD/ACE2 protein complex. These considerations make us hypothesize that an increase of less than 5 % in the calculated interaction energies between the VoC-RBDs and ACE2, compared to interactions energies calculated for the Wuhan spike RBD/ACE2 protein complex, should make still sensitive the investigated VoC to the current employed Wuhan SARS-CoV-2 based vaccines [63, 64], although the slightly increased VoC-RBD/ACE2 binding interactions might determine a lower affinity for the vaccine induced antibodies [12–14]. Conversely, VoC showing more than 3 mutations on the spike RBD head and/or an increase of more than 5-10% in the calculated interaction energies with ACE2 should deserve great attention. In this context interaction energies calculated by using the described molecular framework can be used for leading/driving *in vitro* binding assays, for estimating the putative aggressiveness of new VoC based on the calculated binding affinities, and for the development of new vaccines and antibodies.

Furthermore, while trying to understand if there was an underestimation of SARS-CoV-2 pre-existing immunity [65], our molecular framework can be used for investigating the resistance shown by some individuals to the infection or to the development of clinical manifestations despite infection, by evaluating the importance of single gene variants in the cell entry factors [24] shown by resistant individuals, starting from putative ACE2 variants [66, 67].

In the end, we wondered about the possible existence of ACE2 structurally related receptors that might interact with SARS-CoV-2 spike RBD in those tissues showing ACE2 poorly expressed [24, 25]. Our fold-recognition based analysis revealed that beyond the expected ACE, THOP1 and NLN appear structurally related to ACE2. ACE, THOP1 and NLN share the “oligopeptidase/receptor” activity with ACE2, and show at least one isoform localized at the plasma membrane [68, 69]. While ACE appears to form a weaker protein complex with the SARS-CoV-2 spike RBD, in our interaction analyses, the interaction energies calculated at the protein-protein interface of the SARS-CoV-2 spike RBD/THOP1 or SARS-CoV-2 spike RBD/NLN show binding energies higher than those calculated for the SARS-CoV-2 spike RBD/ACE2 protein complex, maybe due to a supplementary alpha-helix located at the interface with the SARS-CoV-2 spike RBD in the obtained 3D protein complexes. NLN and THOP1 participate to the cleavage of cytosolic peptides and share the 80% of identical residues [69].

While the expression of NLN or ACE is comparable to or slightly higher than ACE2 expression in all the investigated tissues, it is possible to see that THOP1 is highly more expressed than ACE2 in most of the screened tissues and, most importantly, it is highly expressed in the lung, in the colon, in the esophagus mucosa and in all the brain compartments.

Although THOP1 and ACE2 show a low percentage of identical residues (< 30%), their related biochemical function and high structural similarity (overall RMSD < 4.5 Å) make THOP1 a suggestive/alternative candidate receptor for the SARS-CoV-2 spike protein. In addition, in our GTEx analyses the high expression of THOP1 appears to correlate with high expression levels of the Furin protease, which may participate to spike cleavage allowing pre-/post-fusion conformational changes crucial for host-cell penetration [24, 70], above all in those tissues showing a low ACE2 expression, i.e., in the lung, in the esophagus mucosa or in the brain. Notably, THOP1 expression was observed upregulated in COVID-19 infected patients and in particular in the effector CD8 T-cells of COVID-19 patients at the beginning of the infection [71]. Furthermore, THOP1 appears to play a crucial role in the regulation of MHC I cell-surface expression [72–74]. Due to the shown high expression of THOP1 in several brain compartments and in the effector CD8 T-cells of COVID-19 patients [71], it raises the question about a possible relationship between different SARS-CoV-2 immune responses [75, 76], neurological disorders observed in long covid patients, and THOP1 involvement in MHC-class I regulation in COVID-19 patients [77–79].

## DECLARATIONS

### Availability of data and material

All data generated or analysed data here presented study are included in this published article and its supplementary information files. 3D atomic coordinate of the generated PDF files are available upon request.

### Code availability

All the necessary information and protein accession numbers are provided along the manuscript

### Ethics approval

NA

### Consent to participate

NA

### Consent for publication

NA

### Authors’ contributions

Conceptualization, S.T., M.V., A.D.G. and C.L.P.; Data curation, V.T., L.L., F.P., A.O.; Formal analysis, V.T., F.P., L.L., A.O.; Methodology, V.T., L.L., F.P., and C.L.P.; Supervision, M.V., A.D.G. and C.L.P.; Writing—original draft, V.T. and C.L.P.;Writing—review & editing, V.T., L.L., F.P., A.O., S.T., M.V., A.D.G. and C.L.P.

## Acknowledgements

Authors would like to thank the Italian Association for Mitochondrial Research (www.mitoairm.it) for hosting the webpage (https://www.mitoairm.it/covid19affinities) that will allow interested researchers to run the presented pipeline for their targets. Authors would also like to thank for the IT resources made available by ReCaS, a project funded by the MIUR (Italian Ministry for Education, University and Re-search) in the “PON Ricerca e Competitività 2007–2013-Azione I-Interventi di rafforzamento strutturale” PONa3_00052, Avviso 254/Ric, University of Bari.

## Abbreviations

SARS: Severe acute respiratory syndrome
CoV: Coronavirus
(RBD): Receptor Binding Domain
(ACE2): angiotensin converting enzyme II
(THOP1): thimet oligopeptidase
(NLN): Neurolysin
(VoC): Variants of Concern

