## Supplementary material for "A modular molecular framework for quickly estimating the binding affinity of the spike protein of SARS-CoV-2 variants for ACE2, in presence of mutations at the spike receptor binding domain": Supp. Tab. 1

**Supp. Tab. 1. SARS-CoV-2 spike RBD/ACE2 interactions estimated throug PIC analysis**

### hCoV.19Wuhan.WIV04.2019
#### 6m0j.pdb

**Hydrophobic Interactions within 5 Angstroms**

| Position | Residue | Chain | Position | Residue | Chain |
|---|---|---|---|---|---|
| 28 | PHE | A | 489 | TYR | E |
| 79 | LEU | A | 486 | PHE | E |
| 82 | MET | A | 486 | PHE | E |
| 83 | TYR | A | 486 | PHE | E |

**Protein-Protein disulphide bridges**

**Protein-Protein Main Chain-Main Chain H-Bonds**

| DONOR | | | | ACCEPTOR | | | | PARAMETERS | | | | |
|---|---|---|---|---|---|---|---|---|---|---|---|---|
| POS | CHAIN | RES | ATOM | POS | CHAIN | RES | ATOM | MO | Dd-a | Dh-a | A(d-H-N) | A(a-O=C) |
| 502 | E | GLY | N | 353 | A | LYS | O | - | 2.79 | 1.82 | 165.71 | 157.43 |

Dd-a = Distance Between Donor and Acceptor
Dh-a = Distance Between Hydrogen and Acceptor
A(d-H-N) = Angle Between Donor-H-N
A(a-O=C) = Angle Between Acceptor-O=C
MO = Multiple Occupancy
Note that angles that are undefined are written as 999.99

**Protein-Protein Main Chain-Side Chain Hydrogen Bonds**

| DONOR | | | | ACCEPTOR | | | | PARAMETERS | | | | |
|---|---|---|---|---|---|---|---|---|---|---|---|---|
| POS | CHAIN | RES | ATOM | POS | CHAIN | RES | ATOM | MO | Dd-a | Dh-a | A(d-H-N) | A(a-O=C) |
| 31 | A | LYS | NZ | 490 | E | PHE | O | - | 3.17 | 9.99 | 999.99 | 147.54 |
| 41 | A | TYR | OH | 500 | E | THR | O | - | 3.18 | 9.99 | 999.99 | 80.93 |
| 353 | A | LYS | NZ | 496 | E | GLY | O | - | 2.91 | 9.99 | 999.99 | 113.76 |
| 355 | A | ASP | OD2 | 500 | E | THR | O | 1 | 3.28 | 3.73 | 57.29 | 116.93 |
| 355 | A | ASP | OD2 | 500 | E | THR | O | 2 | 3.28 | 3.27 | 80.76 | 116.93 |
| 501 | E | ASN | N | 41 | A | TYR | OH | - | 3.43 | 3.96 | 51.32 | 999.99 |

Dd-a = Distance Between Donor and Acceptor
Dh-a = Distance Between Hydrogen and Acceptor
A(d-H-N) = Angle Between Donor-H-N
A(a-O=C) = Angle Between Acceptor-O=C
MO = Multiple Occupancy
Note that angles that are undefined are written as 999.99

**Protein-Protein Side Chain-Side Chain Hydrogen Bonds**

| DONOR | | | | ACCEPTOR | | | | PARAMETERS | | | | |
|---|---|---|---|---|---|---|---|---|---|---|---|---|
| POS | CHAIN | RES | ATOM | POS | CHAIN | RES | ATOM | MO | Dd-a | Dh-a | A(d-H-N) | A(a-O=C) |
| 24 | A | GLN | OE1 | 487 | E | ASN | ND2 | 1 | 2.86 | 3.16 | 64.30 | 999.99 |
| 24 | A | GLN | OE1 | 487 | E | ASN | ND2 | 2 | 2.86 | 2.24 | 115.24 | 999.99 |
| 31 | A | LYS | NZ | 493 | E | GLN | OE1 | - | 2.92 | 9.99 | 999.99 | 999.99 |
| 35 | A | GLU | OE2 | 493 | E | GLN | OE1 | 1 | 3.46 | 4.04 | 50.46 | 999.99 |
| 35 | A | GLU | OE2 | 493 | E | GLN | OE1 | 2 | 3.46 | 2.40 | 173.88 | 999.99 |
| 35 | A | GLU | OE2 | 493 | E | GLN | NE2 | 1 | 2.88 | 3.46 | 48.99 | 999.99 |
| 35 | A | GLU | OE2 | 493 | E | GLN | NE2 | 2 | 2.88 | 2.22 | 117.88 | 999.99 |
| 42 | A | GLN | NE2 | 449 | E | TYR | OH | 1 | 2.92 | 3.12 | 69.65 | 999.99 |
| 42 | A | GLN | NE2 | 449 | E | TYR | OH | 2 | 2.92 | 2.14 | 129.15 | 999.99 |
| 83 | A | TYR | OH | 487 | E | ASN | OD1 | - | 2.83 | 9.99 | 999.99 | 999.99 |
| 417 | E | LYS | NZ | 30 | A | ASP | OD2 | - | 2.88 | 9.99 | 999.99 | 999.99 |
| 449 | E | TYR | OH | 38 | A | ASP | OD1 | - | 3.05 | 9.99 | 999.99 | 999.99 |
| 449 | E | TYR | OH | 38 | A | ASP | OD2 | - | 2.80 | 9.99 | 999.99 | 999.99 |
| 449 | E | TYR | OH | 42 | A | GLN | NE2 | - | 2.92 | 9.99 | 999.99 | 999.99 |
| 487 | E | ASN | ND2 | 24 | A | GLN | OE1 | 1 | 2.86 | 1.95 | 143.07 | 999.99 |
| 487 | E | ASN | ND2 | 24 | A | GLN | OE1 | 2 | 2.86 | 3.31 | 56.31 | 999.99 |
| 487 | E | ASN | OD1 | 83 | A | TYR | OH | 1 | 2.83 | 2.38 | 103.99 | 999.99 |
| 487 | E | ASN | OD1 | 83 | A | TYR | OH | 2 | 2.83 | 2.70 | 85.90 | 999.99 |
| 493 | E | GLN | OE1 | 35 | A | GLU | OE2 | 1 | 3.46 | 2.71 | 126.90 | 999.99 |
| 493 | E | GLN | OE1 | 35 | A | GLU | OE2 | 2 | 3.46 | 4.40 | 24.73 | 999.99 |
| 493 | E | GLN | NE2 | 35 | A | GLU | OE2 | 1 | 2.88 | 1.87 | 156.82 | 999.99 |
| 493 | E | GLN | NE2 | 35 | A | GLU | OE2 | 2 | 2.88 | 3.54 | 43.80 | 999.99 |
| 500 | E | THR | OG1 | 41 | A | TYR | OH | - | 2.74 | 9.99 | 999.99 | 999.99 |

Dd-a = Distance Between Donor and Acceptor
Dh-a = Distance Between Hydrogen and Acceptor
A(d-H-N) = Angle Between Donor-H-N
A(a-O=C) = Angle Between Acceptor-O=C
MO = Multiple Occupancy
Note that angles that are undefined are written as 999.99

**Protein-Protein Ionic Interactions within 6 anstroms**

| Position | Residue | Chain | Position | Residue | Chain |
|---|---|---|---|---|---|
| 30 | ASP | A | 417 | LYS | E |
| 31 | LYS | A | 484 | GLU | E |

**Protein-Protein Aromatic-Aromatic Interactions within 4.5 and 7 Angstroms**

| Position | Residue | Chain | Position | Residue | Chain | D(centroid-centroid) | Dihedral Angle |
|---|---|---|---|---|---|---|---|
| 83 | TYR | A | 486 | PHE | E | 4.84 | 151.25 |

**Protein-Protein Aromatic-Sulphur Interactions within 5.3 Angstroms**

| Position | Residue | Chain | Position | Residue | Chain | D(centroid-centroid) | Angle |
|---|---|---|---|---|---|---|---|
| 486 | PHE | E | 82 | MET | A | 3.93 | 146.32 |

**Protein-protein cation-pi interactions**

# B.1.1.7_UK
# S494P_N501Y_E484K

**Hydrophobic Interactions within 5 Angstroms**

| Position | Residue | Chain | Position | Residue | Chain |
|---|---|---|---|---|---|
| 28 | PHE | A | 489 | TYR | E |
| 41 | TYR | A | 501 | TYR | E |
| 79 | LEU | A | 486 | PHE | E |
| 82 | MET | A | 486 | PHE | E |
| 83 | TYR | A | 486 | PHE | E |

**Protein-Protein disulphide bridges**

**Protein-Protein Main Chain-Main Chain H-Bonds**

| DONOR | | | | ACCEPTOR | | | | PARAMETERS | | | | |
|---|---|---|---|---|---|---|---|---|---|---|---|---|
| POS | CHAIN | RES | ATOM | POS | CHAIN | RES | ATOM | MO | Dd-a | Dh-a | A(d-H-N) | A(a-O=C) |
| 502 | E | GLY | N | 353 | A | LYS | O | - | 2.85 | 1.89 | 166.49 | 150.73 |

Dd-a = Distance Between Donor and Acceptor
Dh-a = Distance Between Hydrogen and Acceptor
A(d-H-N) = Angle Between Donor-H-N
A(a-O=C) = Angle Between Acceptor-O=C
MO = Multiple Occupancy
Note that angles that are undefined are written as 999.99

**Protein-Protein Main Chain-Side Chain Hydrogen Bonds**

| DONOR | | | | ACCEPTOR | | | | PARAMETERS | | | | |
|---|---|---|---|---|---|---|---|---|---|---|---|---|
| POS | CHAIN | RES | ATOM | POS | CHAIN | RES | ATOM | MO | Dd-a | Dh-a | A(d-H-N) | A(a-O=C) |
| 355 | A | ASP | OD2 | 500 | E | THR | O | 1 | 3.20 | 3.60 | 60.23 | 115.24 |
| 355 | A | ASP | OD2 | 500 | E | THR | O | 2 | 3.20 | 3.24 | 78.37 | 115.24 |

Dd-a = Distance Between Donor and Acceptor
Dh-a = Distance Between Hydrogen and Acceptor
A(d-H-N) = Angle Between Donor-H-N
A(a-O=C) = Angle Between Acceptor-O=C

MO = Multiple Occupancy
Note that angles that are undefined are written as 999.99

**Protein-Protein Side Chain-Side Chain Hydrogen Bonds**

| DONOR | | | | ACCEPTOR | | | | PARAMETERS | | | | |
|---|---|---|---|---|---|---|---|---|---|---|---|---|
| POS | CHAIN | RES | ATOM | POS | CHAIN | RES | ATOM | MO | Dd-a | Dh-a | A(d-H-N) | A(a-O=C) |
| 24 | A | GLN | OE1 | 487 | E | ASN | ND2 | 1 | 2.86 | 3.15 | 64.81 | 999.99 |
| 24 | A | GLN | OE1 | 487 | E | ASN | ND2 | 2 | 2.86 | 2.30 | 111.12 | 999.99 |
| 31 | A | LYS | NZ | 493 | E | GLN | OE1 | - | 2.88 | 9.99 | 999.99 | 999.99 |
| 31 | A | LYS | NZ | 493 | E | GLN | NE2 | - | 3.43 | 9.99 | 999.99 | 999.99 |
| 83 | A | TYR | OH | 487 | E | ASN | OD1 | - | 2.50 | 9.99 | 999.99 | 999.99 |
| 353 | A | LYS | NZ | 501 | E | TYR | OH | - | 2.90 | 9.99 | 999.99 | 999.99 |
| 417 | E | LYS | NZ | 30 | A | ASP | OD2 | - | 2.59 | 9.99 | 999.99 | 999.99 |
| 449 | E | TYR | OH | 38 | A | ASP | OD1 | - | 2.90 | 9.99 | 999.99 | 999.99 |
| 449 | E | TYR | OH | 38 | A | ASP | OD2 | - | 2.83 | 9.99 | 999.99 | 999.99 |
| 487 | E | ASN | ND2 | 24 | A | GLN | OE1 | 1 | 2.86 | 2.29 | 112.31 | 999.99 |
| 487 | E | ASN | ND2 | 24 | A | GLN | OE1 | 2 | 2.86 | 3.05 | 69.57 | 999.99 |
| 487 | E | ASN | OD1 | 83 | A | TYR | OH | 1 | 2.50 | 2.04 | 102.46 | 999.99 |
| 487 | E | ASN | OD1 | 83 | A | TYR | OH | 2 | 2.50 | 2.36 | 84.59 | 999.99 |
| 500 | E | THR | OG1 | 41 | A | TYR | OH | - | 2.71 | 9.99 | 999.99 | 999.99 |

Dd-a = Distance Between Donor and Acceptor
Dh-a = Distance Between Hydrogen and Acceptor
A(d-H-N) = Angle Between Donor-H-N
A(a-O=C) = Angle Between Acceptor-O=C
MO = Multiple Occupancy
Note that angles that are undefined are written as 999.99

**Protein-Protein Ionic Interactions within 6 anstroms**

| Position | Residue | Chain | Position | Residue | Chain |
|---|---|---|---|---|---|
| 30 | ASP | A | 417 | LYS | E |

**Protein-Protein Aromatic-Aromatic Interactions within 4.5 and 7 Angstroms**

| Position | Residue | Chain | Position | Residue | Chain | D(centroid-centroid) | Dihedral Angle |
|---|---|---|---|---|---|---|---|
| 41 | TYR | A | 501 | TYR | E | 4.74 | 79.53 |
| 83 | TYR | A | 486 | PHE | E | 4.85 | 149.95 |

**Protein-Protein Aromatic-Sulphur Interactions within 5.3 Angstroms**

| Position | Residue | Chain | Position | Residue | Chain | D(centroid-centroid) | Angle |
|---|---|---|---|---|---|---|---|
| 486 | PHE | E | 82 | MET | A | 3.85 | 146.23 |

**Protein-protein cation-pi interactions**

| | | | | | | | |
|---|---|---|---|---|---|---|---|
| 501 | TYR | E | 353 | LYS | A | 4.53 | 55.81 |

### B.1.315_S.africa
# N501Y_E484K_K417T

**Hydrophobic Interactions within 5 Angstroms**

| Position | Residue | Chain | Position | Residue | Chain |
|---|---|---|---|---|---|
| 28 | PHE | A | 489 | TYR | E |
| 41 | TYR | A | 501 | TYR | E |
| 79 | LEU | A | 486 | PHE | E |
| 82 | MET | A | 486 | PHE | E |
| 83 | TYR | A | 486 | PHE | E |

**Protein-Protein disulphide bridges**

**Protein-Protein Main Chain-Main Chain H-Bonds**

| DONOR | | | | ACCEPTOR | | | | PARAMETERS | | | | |
|---|---|---|---|---|---|---|---|---|---|---|---|---|
| POS | CHAIN | RES | ATOM | POS | CHAIN | RES | ATOM | MO | Dd-a | Dh-a | A(d-H-N) | A(a-O=C) |
| 502 | E | GLY | N | 353 | A | LYS | O | - | 2.80 | 1.84 | 167.07 | 152.29 |

Dd-a = Distance Between Donor and Acceptor
Dh-a = Distance Between Hydrogen and Acceptor
A(d-H-N) = Angle Between Donor-H-N
A(a-O=C) = Angle Between Acceptor-O=C
MO = Multiple Occupancy
Note that angles that are undefined are written as 999.99

**Protein-Protein Main Chain-Side Chain Hydrogen Bonds**

| DONOR | | | | ACCEPTOR | | | | PARAMETERS | | | | |
|---|---|---|---|---|---|---|---|---|---|---|---|---|
| POS | CHAIN | RES | ATOM | POS | CHAIN | RES | ATOM | MO | Dd-a | Dh-a | A(d-H-N) | A(a-O=C) |
| 28 | A | PHE | N | 489 | E | TYR | OH | - | 3.34 | 3.67 | 63.16 | 999.99 |
| 31 | A | LYS | NZ | 490 | E | PHE | O | - | 2.94 | 9.99 | 999.99 | 149.04 |
| 31 | A | LYS | NZ | 492 | E | LEU | O | - | 3.26 | 9.99 | 999.99 | 126.57 |
| 41 | A | TYR | OH | 500 | E | THR | O | - | 3.45 | 9.99 | 999.99 | 70.82 |
| 355 | A | ASP | OD2 | 500 | E | THR | O | 1 | 3.36 | 3.83 | 56.64 | 114.32 |
| 355 | A | ASP | OD2 | 500 | E | THR | O | 2 | 3.36 | 3.34 | 82.13 | 114.32 |

| | | | | | | | | | | |
|---|---|---|---|---|---|---|---|---|---|---|
| 501 E | TYR | N | 41 A | TYR | OH | - | 3.34 | 3.74 | 59.11 | 999.99 |

Dd-a = Distance Between Donor and Acceptor
Dh-a = Distance Between Hydrogen and Acceptor
A(d-H-N) = Angle Between Donor-H-N
A(a-O=C) = Angle Between Acceptor-O=C
MO = Multiple Occupancy
Note that angles that are undefined are written as 999.99

**Protein-Protein Side Chain-Side Chain Hydrogen Bonds**

| DONOR | | | ACCEPTOR | | | PARAMETERS | | | | |
|---|---|---|---|---|---|---|---|---|---|---|
| 24 A | GLN | OE1 | 487 E | ASN | ND2 | 1 | 2.89 | 3.06 | 71.24 | 999.99 |
| 24 A | GLN | OE1 | 487 E | ASN | ND2 | 2 | 2.89 | 2.39 | 107.49 | 999.99 |
| 31 A | LYS | NZ | 493 E | GLN | OE1 | - | 2.88 | 9.99 | 999.99 | 999.99 |
| 35 A | GLU | OE1 | 493 E | GLN | NE2 | 1 | 2.93 | 2.72 | 90.88 | 999.99 |
| 35 A | GLU | OE1 | 493 E | GLN | NE2 | 2 | 2.93 | 2.34 | 113.35 | 999.99 |
| 42 A | GLN | NE2 | 449 E | TYR | OH | 1 | 3.49 | 3.89 | 60.23 | 999.99 |
| 42 A | GLN | NE2 | 449 E | TYR | OH | 2 | 3.49 | 2.86 | 118.58 | 999.99 |
| 83 A | TYR | OH | 487 E | ASN | OD1 | - | 2.57 | 9.99 | 999.99 | 999.99 |
| 353 A | LYS | NZ | 501 E | TYR | OH | - | 2.92 | 9.99 | 999.99 | 999.99 |
| 449 E | TYR | OH | 38 A | ASP | OD1 | - | 2.89 | 9.99 | 999.99 | 999.99 |
| 449 E | TYR | OH | 38 A | ASP | OD2 | - | 2.81 | 9.99 | 999.99 | 999.99 |
| 449 E | TYR | OH | 42 A | GLN | NE2 | - | 3.49 | 9.99 | 999.99 | 999.99 |
| 487 E | ASN | ND2 | 24 A | GLN | OE1 | 1 | 2.89 | 2.36 | 109.73 | 999.99 |
| 487 E | ASN | ND2 | 24 A | GLN | OE1 | 2 | 2.89 | 3.30 | 58.69 | 999.99 |
| 487 E | ASN | OD1 | 83 A | TYR | OH | 1 | 2.57 | 2.16 | 99.51 | 999.99 |
| 487 E | ASN | OD1 | 83 A | TYR | OH | 2 | 2.57 | 2.36 | 88.96 | 999.99 |
| 493 E | GLN | NE2 | 35 A | GLU | OE1 | 1 | 2.93 | 1.90 | 164.84 | 999.99 |
| 493 E | GLN | NE2 | 35 A | GLU | OE1 | 2 | 2.93 | 3.37 | 56.79 | 999.99 |
| 500 E | THR | OG1 | 41 A | TYR | OH | - | 2.58 | 9.99 | 999.99 | 999.99 |

Dd-a = Distance Between Donor and Acceptor
Dh-a = Distance Between Hydrogen and Acceptor
A(d-H-N) = Angle Between Donor-H-N
A(a-O=C) = Angle Between Acceptor-O=C

MO = Multiple Occupancy
Note that angles that are undefined are written as 999.99

**Protein-Protein Ionic Interactions within 6 anstroms**

| Position | Residue | Chain | Position | Residue | Chain |
|---|---|---|---|---|---|

**Protein-Protein Aromatic-Aromatic Interactions within 4.5 and 7 Angstroms**

| Position | Residue | Chain | Position | Residue | Chain | D(centroid-c | Dihedral Angle |
|---|---|---|---|---|---|---|---|
| 41 | TYR | A | 501 | TYR | E | 4.75 | 69.83 |
| 83 | TYR | A | 486 | PHE | E | 4.86 | 145.32 |

**Protein-Protein Aromatic-Sulphur Interactions within 5.3 Angstroms**

| Position | Residue | Chain | Position | Residue | Chain | D(centroid-c | Angle |
|---|---|---|---|---|---|---|---|
| 486 | PHE | E | 82 | MET | A | 3.89 | 158.30 |

**Protein-protein cation-pi interactions within 6 angstrom**

| | | | | | | | |
|---|---|---|---|---|---|---|---|
| 501 | TYR | E | 353 | LYS | A | 4.64 | 59.07 |

### B.1.427 California
# L452R

**Hydrophobic Interactions within 5 Angstroms**

| Position | Residue | Chain | Position | Residue | Chain |
|---|---|---|---|---|---|
| 28 | PHE | A | 489 | TYR | E |
| 79 | LEU | A | 486 | PHE | E |
| 82 | MET | A | 486 | PHE | E |
| 83 | TYR | A | 486 | PHE | E |

**Protein-Protein disulphide bridges**

**Protein-Protein Main Chain-Main Chain H-Bonds**

| DONOR | | | | ACCEPTOR | | | | PARAMETERS | | | | |
|---|---|---|---|---|---|---|---|---|---|---|---|---|
| POS | CHAIN | RES | ATOM | POS | CHAIN | RES | ATOM | MO | Dd-a | Dh-a | A(d-H-N) | A(a-O=C) |
| 502 | E | GLY | N | 353 | A | LYS | O | - | 2.76 | 1.80 | 165.44 | 154.27 |

Dd-a = Distance Between Donor and Acceptor
Dh-a = Distance Between Hydrogen and Acceptor
A(d-H-N) = Angle Between Donor-H-N
A(a-O=C) = Angle Between Acceptor-O=C
MO = Multiple Occupancy
Note that angles that are undefined are written as 999.99

**Protein-Protein Main Chain-Side Chain Hydrogen Bonds**

| DONOR | | | | ACCEPTOR | | | | PARAMETERS | | | | |
|---|---|---|---|---|---|---|---|---|---|---|---|---|
| POS | CHAIN | RES | ATOM | POS | CHAIN | RES | ATOM | MO | Dd-a | Dh-a | A(d-H-N) | A(a-O=C) |
| 41 | A | TYR | OH | 500 | E | THR | O | - | 3.31 | 9.99 | 999.99 | 77.37 |
| 353 | A | LYS | NZ | 496 | E | GLY | O | - | 2.89 | 9.99 | 999.99 | 115.89 |
| 355 | A | ASP | OD2 | 500 | E | THR | O | 1 | 3.45 | 3.96 | 54.26 | 113.81 |
| 355 | A | ASP | OD2 | 500 | E | THR | O | 2 | 3.45 | 3.46 | 80.72 | 113.81 |
| 501 | E | ASN | N | 41 | A | TYR | OH | - | 3.47 | 3.91 | 56.78 | 999.99 |

Dd-a = Distance Between Donor and Acceptor
Dh-a = Distance Between Hydrogen and Acceptor
A(d-H-N) = Angle Between Donor-H-N
A(a-O=C) = Angle Between Acceptor-O=C
MO = Multiple Occupancy
Note that angles that are undefined are written as 999.99

**Protein-Protein Side Chain-Side Chain Hydrogen Bonds**

| DONOR | | | | ACCEPTOR | | | | PARAMETERS | | | | |
|---|---|---|---|---|---|---|---|---|---|---|---|---|
| POS | CHAIN | RES | ATOM | POS | CHAIN | RES | ATOM | MO | Dd-a | Dh-a | A(d-H-N) | A(a-O=C) |
| 24 | A | GLN | OE1 | 487 | E | ASN | ND2 | 1 | 2.99 | 3.19 | 69.63 | 999.99 |
| 24 | A | GLN | OE1 | 487 | E | ASN | ND2 | 2 | 2.99 | 2.07 | 142.89 | 999.99 |
| 31 | A | LYS | NZ | 493 | E | GLN | OE1 | - | 2.90 | 9.99 | 999.99 | 999.99 |
| 35 | A | GLU | OE2 | 493 | E | GLN | NE2 | 1 | 2.87 | 3.30 | 57.73 | 999.99 |
| 35 | A | GLU | OE2 | 493 | E | GLN | NE2 | 2 | 2.87 | 2.15 | 122.78 | 999.99 |
| 42 | A | GLN | NE2 | 449 | E | TYR | OH | 1 | 3.08 | 3.39 | 64.06 | 999.99 |
| 42 | A | GLN | NE2 | 449 | E | TYR | OH | 2 | 3.08 | 2.53 | 111.93 | 999.99 |
| 83 | A | TYR | OH | 487 | E | ASN | OD1 | - | 3.11 | 9.99 | 999.99 | 999.99 |
| 83 | A | TYR | OH | 487 | E | ASN | ND2 | - | 3.27 | 9.99 | 999.99 | 999.99 |
| 417 | E | LYS | NZ | 30 | A | ASP | OD2 | - | 2.62 | 9.99 | 999.99 | 999.99 |
| 449 | E | TYR | OH | 38 | A | ASP | OD1 | - | 3.09 | 9.99 | 999.99 | 999.99 |
| 449 | E | TYR | OH | 38 | A | ASP | OD2 | - | 2.80 | 9.99 | 999.99 | 999.99 |
| 449 | E | TYR | OH | 42 | A | GLN | NE2 | - | 3.08 | 9.99 | 999.99 | 999.99 |
| 487 | E | ASN | ND2 | 24 | A | GLN | OE1 | 1 | 2.99 | 2.53 | 105.19 | 999.99 |
| 487 | E | ASN | ND2 | 24 | A | GLN | OE1 | 2 | 2.99 | 3.47 | 54.51 | 999.99 |
| 487 | E | ASN | OD1 | 83 | A | TYR | OH | 1 | 3.11 | 2.75 | 99.12 | 999.99 |
| 487 | E | ASN | OD1 | 83 | A | TYR | OH | 2 | 3.11 | 3.73 | 47.31 | 999.99 |
| 487 | E | ASN | ND2 | 83 | A | TYR | OH | 1 | 3.27 | 2.95 | 98.41 | 999.99 |
| 487 | E | ASN | ND2 | 83 | A | TYR | OH | 2 | 3.27 | 3.97 | 42.82 | 999.99 |
| 493 | E | GLN | NE2 | 35 | A | GLU | OE2 | 1 | 2.87 | 1.82 | 178.96 | 999.99 |
| 493 | E | GLN | NE2 | 35 | A | GLU | OE2 | 2 | 2.87 | 3.46 | 48.35 | 999.99 |
| 500 | E | THR | OG1 | 41 | A | TYR | OH | - | 2.70 | 9.99 | 999.99 | 999.99 |
| 505 | E | TYR | OH | 37 | A | GLU | OE1 | - | 2.77 | 9.99 | 999.99 | 999.99 |
| 505 | E | TYR | OH | 37 | A | GLU | OE2 | - | 3.28 | 9.99 | 999.99 | 999.99 |

Dd-a = Distance Between Donor and Acceptor
Dh-a = Distance Between Hydrogen and Acceptor
A(d-H-N) = Angle Between Donor-H-N
A(a-O=C) = Angle Between Acceptor-O=C
MO = Multiple Occupancy
Note that angles that are undefined are written as 999.99

**Protein-Protein Ionic Interactions within 6 anstroms**

| Position | Residue | Chain | Position | Residue | Chain |
|---|---|---|---|---|---|
| 30 | ASP | A | 417 | LYS | E |
| 31 | LYS | A | 484 | GLU | E |
| 37 | GLU | A | 403 | ARG | E |

**Protein-Protein Aromatic-Aromatic Interactions within 4.5 and 7 Angstroms**

| Position | Residue | Chain | Position | Residue | Chain | D(centroid- | Dihedral Angle |
|---|---|---|---|---|---|---|---|
| 83 | TYR | A | 486 | PHE | E | 4.99 | 143.57 |

**Protein-Protein Aromatic-Sulphur Interactions within 5.3 Angstroms**

| Position | Residue | Chain | Position | Residue | Chain | D(centroid- | Angle |
|---|---|---|---|---|---|---|---|
| 486 | PHE | E | 82 | MET | A | 3.89 | 146.68 |

**Protein-protein cation-pi interactions**

### B1.617.india
# E484Q-L452R

**Hydrophobic Interactions within 5 Angstroms**

| Position | Residue | Chain | Position | Residue | Chain |
|---|---|---|---|---|---|
| 28 | PHE | A | 489 | TYR | E |
| 79 | LEU | A | 486 | PHE | E |
| 82 | MET | A | 486 | PHE | E |
| 83 | TYR | A | 486 | PHE | E |

**Protein-Protein disulphide bridges**

**Protein-Protein Main Chain-Main Chain H-Bonds**

| DONOR | | | | ACCEPTOR | | | | PARAMETERS | | | | |
|---|---|---|---|---|---|---|---|---|---|---|---|---|
| POS | CHAIN | RES | ATOM | POS | CHAIN | RES | ATOM | MO | Dd-a | Dh-a | A(d-H-N) | A(a-O=C) |
| 502 | E | GLY | N | 353 | A | LYS | O | - | 2.79 | 1.82 | 165.71 | 157.43 |

Dd-a = Distance Between Donor and Acceptor
Dh-a = Distance Between Hydrogen and Acceptor
A(d-H-N) = Angle Between Donor-H-N
A(a-O=C) = Angle Between Acceptor-O=C
MO = Multiple Occupancy
Note that angles that are undefined are written as 999.99

**Protein-Protein Main Chain-Side Chain Hydrogen Bonds**

| DONOR | | | | ACCEPTOR | | | | PARAMETERS | | | | |
|---|---|---|---|---|---|---|---|---|---|---|---|---|
| POS | CHAIN | RES | ATOM | POS | CHAIN | RES | ATOM | MO | Dd-a | Dh-a | A(d-H-N) | A(a-O=C) |
| 353 | A | LYS | NZ | 496 | E | GLY | O | - | 2.92 | 9.99 | 999.99 | 114.62 |
| 355 | A | ASP | OD2 | 500 | E | THR | O | 1 | 3.20 | 3.57 | 61.06 | 112.64 |
| 355 | A | ASP | OD2 | 500 | E | THR | O | 2 | 3.20 | 3.23 | 78.78 | 112.64 |

Dd-a = Distance Between Donor and Acceptor
Dh-a = Distance Between Hydrogen and Acceptor
A(d-H-N) = Angle Between Donor-H-N
A(a-O=C) = Angle Between Acceptor-O=C
MO = Multiple Occupancy
Note that angles that are undefined are written as 999.99

**Protein-Protein Side Chain-Side Chain Hydrogen Bonds**

| DONOR | | | | ACCEPTOR | | | | PARAMETERS | | | | |
|---|---|---|---|---|---|---|---|---|---|---|---|---|
| POS | CHAIN | RES | ATOM | POS | CHAIN | RES | ATOM | MO | Dd-a | Dh-a | A(d-H-N) | A(a-O=C) |
| 24 | A | GLN | OE1 | 487 | E | ASN | ND2 | 1 | 2.70 | 2.84 | 71.51 | 999.99 |
| 24 | A | GLN | OE1 | 487 | E | ASN | ND2 | 2 | 2.70 | 2.22 | 104.88 | 999.99 |
| 31 | A | LYS | NZ | 493 | E | GLN | OE1 | - | 2.66 | 9.99 | 999.99 | 999.99 |
| 35 | A | GLU | OE2 | 493 | E | GLN | NE2 | 1 | 2.92 | 3.48 | 50.50 | 999.99 |
| 35 | A | GLU | OE2 | 493 | E | GLN | NE2 | 2 | 2.92 | 2.19 | 123.47 | 999.99 |
| 42 | A | GLN | NE2 | 449 | E | TYR | OH | 1 | 3.26 | 3.52 | 66.89 | 999.99 |
| 42 | A | GLN | NE2 | 449 | E | TYR | OH | 2 | 3.26 | 3.02 | 93.71 | 999.99 |
| 83 | A | TYR | OH | 487 | E | ASN | OD1 | - | 2.58 | 9.99 | 999.99 | 999.99 |
| 449 | E | TYR | OH | 38 | A | ASP | OD1 | - | 2.98 | 9.99 | 999.99 | 999.99 |
| 449 | E | TYR | OH | 38 | A | ASP | OD2 | - | 2.84 | 9.99 | 999.99 | 999.99 |
| 449 | E | TYR | OH | 42 | A | GLN | NE2 | - | 3.26 | 9.99 | 999.99 | 999.99 |
| 487 | E | ASN | ND2 | 24 | A | GLN | OE1 | 1 | 2.70 | 2.10 | 113.90 | 999.99 |
| 487 | E | ASN | ND2 | 24 | A | GLN | OE1 | 2 | 2.70 | 2.92 | 67.65 | 999.99 |
| 487 | E | ASN | OD1 | 83 | A | TYR | OH | 1 | 2.58 | 2.20 | 98.07 | 999.99 |
| 487 | E | ASN | OD1 | 83 | A | TYR | OH | 2 | 2.58 | 2.28 | 93.61 | 999.99 |
| 493 | E | GLN | NE2 | 35 | A | GLU | OE2 | 1 | 2.92 | 1.95 | 150.48 | 999.99 |
| 493 | E | GLN | NE2 | 35 | A | GLU | OE2 | 2 | 2.92 | 3.54 | 46.46 | 999.99 |
| 500 | E | THR | OG1 | 41 | A | TYR | OH | - | 2.49 | 9.99 | 999.99 | 999.99 |

Dd-a = Distance Between Donor and Acceptor

Dh-a = Distance Between Hydrogen and Acceptor
A(d-H-N) = Angle Between Donor-H-N
A(a-O=C) = Angle Between Acceptor-O=C
MO = Multiple Occupancy
Note that angles that are undefined are written as 999.99

**Protein-Protein Ionic Interactions within 6 anstroms**

| Position | Residue | Chain | Position | Residue | Chain |
|---|---|---|---|---|---|
| 30 | ASP | A | 417 | LYS | E |

**Protein-Protein Aromatic-Aromatic Interactions within 4.5 and 7 Angstroms**

| Position | Residue | Chain | Position | Residue | Chain | D(centroic | Dihedral Angle |
|---|---|---|---|---|---|---|---|
| 83 | TYR | A | 486 | PHE | E | 4.88 | 150.76 |

**Protein-Protein Aromatic-Sulphur Interactions within 5.3 Angstroms**

| Position | Residue | Chain | Position | Residue | Chain | D(centroic | Angle |
|---|---|---|---|---|---|---|---|
| 486 | PHE | E | 82 | MET | A | 3.96 | 147.74 |

**Protein-protein cation-pi interactions**

### P.1. Japan Brazil
# N501Y_E484K_K417N

**Hydrophobic Interactions within 5 Angstroms**

| Position | Residue | Chain | Position | Residue | Chain |
|---|---|---|---|---|---|
| 28 | PHE | A | 489 | TYR | E |
| 41 | TYR | A | 501 | TYR | E |
| 79 | LEU | A | 486 | PHE | E |
| 82 | MET | A | 486 | PHE | E |
| 83 | TYR | A | 486 | PHE | E |

**Protein-Protein disulphide bridges**

**Protein-Protein Main Chain-Main Chain H-Bonds**

| DONOR | | | | ACCEPTOR | | | | PARAMETERS | | | | |
|---|---|---|---|---|---|---|---|---|---|---|---|---|
| POS | CHAIN | RES | ATOM | POS | CHAIN | RES | ATOM | MO | Dd-a | Dh-a | A(d-H-N) | A(a-O=C) |
| 502 | E | GLY | N | 353 | A | LYS | O | - | 2.87 | 1.92 | 164.06 | 146.67 |

Dd-a = Distance Between Donor and Acceptor
Dh-a = Distance Between Hydrogen and Acceptor
A(d-H-N) = Angle Between Donor-H-N
A(a-O=C) = Angle Between Acceptor-O=C
MO = Multiple Occupancy
Note that angles that are undefined are written as 999.99

**Protein-Protein Main Chain-Side Chain Hydrogen Bonds**

| DONOR | | | | ACCEPTOR | | | | PARAMETERS | | | | |
|---|---|---|---|---|---|---|---|---|---|---|---|---|
| POS | CHAIN | RES | ATOM | POS | CHAIN | RES | ATOM | MO | Dd-a | Dh-a | A(d-H-N) | A(a-O=C) |
| 28 | A | PHE | N | 489 | E | TYR | OH | - | 3.37 | 3.73 | 61.55 | 999.99 |
| 31 | A | LYS | NZ | 490 | E | PHE | O | - | 3.06 | 9.99 | 999.99 | 158.14 |

| | | | | | | | | | | | | |
|---|---|---|---|---|---|---|---|---|---|---|---|---|
| 353 | A | LYS | NZ | 496 | E | GLY | O | - | 3.23 | 9.99 | 999.99 | 100.75 |
| 355 | A | ASP | OD2 | 500 | E | THR | O | 1 | 3.24 | 3.48 | 67.95 | 112.22 |
| 355 | A | ASP | OD2 | 500 | E | THR | O | 2 | 3.24 | 3.50 | 66.78 | 112.22 |

Dd-a = Distance Between Donor and Acceptor
Dh-a = Distance Between Hydrogen and Acceptor
A(d-H-N) = Angle Between Donor-H-N
A(a-O=C) = Angle Between Acceptor-O=C
MO = Multiple Occupancy
Note that angles that are undefined are written as 999.99

**Protein-Protein Side Chain-Side Chain Hydrogen Bonds**

| DONOR | | | | ACCEPTOR | | | | PARAMETERS | | | | |
|---|---|---|---|---|---|---|---|---|---|---|---|---|
| POS | CHAIN | RES | ATOM | POS | CHAIN | RES | ATOM | MO | Dd-a | Dh-a | A(d-H-N) | A(a-O=C) |
| 24 | A | GLN | OE1 | 487 | E | ASN | ND2 | 1 | 2.85 | 3.10 | 66.52 | 999.99 |
| 24 | A | GLN | OE1 | 487 | E | ASN | ND2 | 2 | 2.85 | 2.27 | 112.22 | 999.99 |
| 31 | A | LYS | NZ | 493 | E | GLN | OE1 | - | 2.92 | 9.99 | 999.99 | 999.99 |
| 35 | A | GLU | OE2 | 493 | E | GLN | OE1 | 1 | 3.13 | 3.54 | 58.67 | 999.99 |
| 35 | A | GLU | OE2 | 493 | E | GLN | OE1 | 2 | 3.13 | 2.10 | 158.44 | 999.99 |
| 35 | A | GLU | OE2 | 493 | E | GLN | NE2 | 1 | 2.91 | 3.26 | 61.68 | 999.99 |
| 35 | A | GLU | OE2 | 493 | E | GLN | NE2 | 2 | 2.91 | 2.08 | 131.42 | 999.99 |
| 42 | A | GLN | NE2 | 449 | E | TYR | OH | 1 | 3.04 | 3.47 | 57.32 | 999.99 |

| | | | | | | | | | | | | |
|---|---|---|---|---|---|---|---|---|---|---|---|---|
| 42 | A | GLN | NE2 | 449 | E | TYR | OH | 2 | 3.04 | 2.02 | 163.61 | 999.99 |
| 83 | A | TYR | OH | 487 | E | ASN | OD1 | - | 2.49 | 9.99 | 999.99 | 999.99 |
| 353 | A | LYS | NZ | 501 | E | TYR | OH | - | 2.91 | 9.99 | 999.99 | 999.99 |
| 449 | E | TYR | OH | 38 | A | ASP | OD1 | - | 2.51 | 9.99 | 999.99 | 999.99 |
| 449 | E | TYR | OH | 42 | A | GLN | NE2 | - | 3.04 | 9.99 | 999.99 | 999.99 |
| 487 | E | ASN | ND2 | 24 | A | GLN | OE1 | 1 | 2.85 | 2.29 | 111.15 | 999.99 |
| 487 | E | ASN | ND2 | 24 | A | GLN | OE1 | 2 | 2.85 | 3.04 | 69.23 | 999.99 |
| 487 | E | ASN | OD1 | 83 | A | TYR | OH | 1 | 2.49 | 1.99 | 104.41 | 999.99 |
| 487 | E | ASN | OD1 | 83 | A | TYR | OH | 2 | 2.49 | 2.26 | 89.43 | 999.99 |
| 493 | E | GLN | OE1 | 35 | A | GLU | OE2 | 1 | 3.13 | 2.26 | 136.40 | 999.99 |
| 493 | E | GLN | OE1 | 35 | A | GLU | OE2 | 2 | 3.13 | 4.01 | 29.23 | 999.99 |
| 493 | E | GLN | NE2 | 35 | A | GLU | OE2 | 1 | 2.91 | 1.94 | 150.71 | 999.99 |
| 493 | E | GLN | NE2 | 35 | A | GLU | OE2 | 2 | 2.91 | 3.70 | 35.01 | 999.99 |
| 500 | E | THR | OG1 | 41 | A | TYR | OH | - | 2.81 | 9.99 | 999.99 | 999.99 |

Dd-a = Distance Between Donor and Acceptor
Dh-a = Distance Between Hydrogen and Acceptor
A(d-H-N) = Angle Between Donor-H-N
A(a-O=C) = Angle Between Acceptor-O=C
MO = Multiple Occupancy

Note that angles that are undefined are written as 999.99

**Protein-Protein Ionic Interactions within 6 anstroms**

| Position | Residue | Chain | Position | Residue | Chain |
|---|---|---|---|---|---|

**Protein-Protein Aromatic-Aromatic Interactions within 4.5 and 7 Angstroms**

| Position | Residue | Chain | Position | Residue | Chain | D(cei | Dihedral Angle |
|---|---|---|---|---|---|---|---|
| 41 | TYR | A | 501 | TYR | E | 4.94 | 78.41 |
| 83 | TYR | A | 486 | PHE | E | 4.98 | 148.19 |

**Protein-Protein Aromatic-Sulphur Interactions within 5.3 Angstroms**

| Position | Residue | Chain | Position | Residue | Chain | D(cei | Angle |
|---|---|---|---|---|---|---|---|
| 486 | PHE | E | 82 | MET | A | 3.79 | 149.78 |

**Protein-protein cation-pi interactions**

| | | | | | | | |
|---|---|---|---|---|---|---|---|
| 501 | TYR | E | 353 | LYS | A | 4.38 | 61.71 |

# B.1.620
# S477N.E484K

**Hydrophobic Interactions within 5 Angstroms**

| Position | Residue | Chain | Position | Residue | Chain |
|---|---|---|---|---|---|
| 28 | PHE | A | 489 | TYR | E |
| 79 | LEU | A | 486 | PHE | E |
| 82 | MET | A | 486 | PHE | E |
| 83 | TYR | A | 486 | PHE | E |

**Protein-Protein disulphide bridges**

NO PROTEIN-PROTEIN DISULPHIDE BRIDGES FOUND

**Protein-Protein Main Chain-Main Chain Hydrogen Bonds**

| DONOR | | | | ACCEPTOR | | | | PARAMETERS | | | | |
|---|---|---|---|---|---|---|---|---|---|---|---|---|
| POS | CHAIN | RES | ATOM | POS | CHAIN | RES | ATOM | MO | Dd-a | Dh-a | A(d-H-N) | A(a-O=C) |
| 502 | E | GLY | N | 353 | A | LYS | O | - | 0,138889 | 0,099306 | 167:12:00 | 153:07:00 |

Dd-a = Distance Between Donor and Acceptor
Dh-a = Distance Between Hydrogen and Acceptor
A(d-H-N) = Angle Between Donor-H-N
A(a-O=C) = Angle Between Acceptor-O=C
MO = Multiple Occupancy
Note that angles that are undefined are written as 999.99

**Protein-Protein Main Chain-Side Chain Hydrogen Bonds**

| DONOR | | | | ACCEPTOR | | | | PARAMETERS | | | | |
|---|---|---|---|---|---|---|---|---|---|---|---|---|
| POS | CHAIN | RES | ATOM | POS | CHAIN | RES | ATOM | MO | Dd-a | Dh-a | A(d-H-N) | A(a-O=C) |
| 41 | A | TYR | OH | 500 | E | THR | O | - | 03:46 | 0,44375 | 999.99 | 74:35:00 |
| 353 | A | LYS | NZ | 496 | E | GLY | O | - | 0,145833 | 0,44375 | 999.99 | 121:58:00 |
| 355 | A | ASP | OD2 | 500 | E | THR | O | 1 | 03:21 | 0,166667 | 60:38:00 | 113.88 |
| 355 | A | ASP | OD2 | 500 | E | THR | O | 2 | 03:21 | 03:37 | 72:50:00 | 113.88 |

Dd-a = Distance Between Donor and Acceptor
Dh-a = Distance Between Hydrogen and Acceptor
A(d-H-N) = Angle Between Donor-H-N
A(a-O=C) = Angle Between Acceptor-O=C
MO = Multiple Occupancy
Note that angles that are undefined are written as 999.99

**Protein-Protein Side Chain-Side Chain Hydrogen Bonds**

| DONOR | | | | ACCEPTOR | | | | PARAMETERS | | | | |
|---|---|---|---|---|---|---|---|---|---|---|---|---|
| POS | CHAIN | RES | ATOM | POS | CHAIN | RES | ATOM | MO | Dd-a | Dh-a | A(d-H-N) | A(a-O=C) |
| 24 | A | GLN | OE1 | 487 | E | ASN | ND2 | 1 | 02:58 | 0,132639 | 71.66 | 999.99 |
| 24 | A | GLN | OE1 | 487 | E | ASN | ND2 | 2 | 02:58 | 02:23 | 96:42:00 | 999.99 |
| 31 | A | LYS | NZ | 493 | E | GLN | OE1 | - | 0,141667 | 0,44375 | 999.99 | 999.99 |
| 35 | A | GLU | OE1 | 493 | E | GLN | OE1 | 1 | 03:28 | 03:44 | 72:06:00 | 999.99 |
| 35 | A | GLU | OE1 | 493 | E | GLN | OE1 | 2 | 03:28 | 0,166667 | 64:22:00 | 999.99 |
| 35 | A | GLU | OE1 | 493 | E | GLN | NE2 | 1 | 0,136111 | 03:09 | 61.99 | 999.99 |
| 35 | A | GLU | OE1 | 493 | E | GLN | NE2 | 2 | 0,136111 | 02:36 | 100:05:00 | 999.99 |
| 42 | A | GLN | NE2 | 449 | E | TYR | OH | 1 | 03:23 | 03:35 | 74:19:00 | 999.99 |

| | | | | | | | | | | | | | |
|---|---|---|---|---|---|---|---|---|---|---|---|---|---|
| 42 | A | GLN | NE2 | 449 | E | TYR | OH | | 2 | 03:23 | 02:52 | 123.71 | 999.99 |
| 83 | A | TYR | OH | 487 | E | ASN | OD1 | - | | 0,136806 | 0,44375 | 999.99 | 999.99 |
| 417 | E | LYS | NZ | 30 | A | ASP | OD2 | - | | 0,142361 | 0,44375 | 999.99 | 999.99 |
| 449 | E | TYR | OH | 38 | A | ASP | OD1 | - | | 03:05 | 0,44375 | 999.99 | 999.99 |
| 449 | E | TYR | OH | 38 | A | ASP | OD2 | - | | 0,128472 | 0,44375 | 999.99 | 999.99 |
| 449 | E | TYR | OH | 42 | A | GLN | NE2 | - | | 03:23 | 0,44375 | 999.99 | 999.99 |
| 487 | E | ASN | ND2 | 24 | A | GLN | OE1 | | 1 | 02:58 | 0,092361 | 133:46:00 | 999.99 |
| 487 | E | ASN | ND2 | 24 | A | GLN | OE1 | | 2 | 02:58 | 03:02 | 55.78 | 999.99 |
| 487 | E | ASN | OD1 | 83 | A | TYR | OH | | 1 | 0,136806 | 02:33 | 102:21:00 | 999.99 |
| 487 | E | ASN | OD1 | 83 | A | TYR | OH | | 2 | 0,136806 | 0,135417 | 79.88 | 999.99 |
| 493 | E | GLN | OE1 | 35 | A | GLU | OE1 | | 1 | 03:28 | 02:49 | 129:24:00 | 999.99 |
| 493 | E | GLN | OE1 | 35 | A | GLU | OE1 | | 2 | 03:28 | 04:22 | 1,020139 | 999.99 |
| 493 | E | GLN | NE2 | 35 | A | GLU | OE1 | | 1 | 0,136111 | 0,092361 | 162.71 | 999.99 |
| 493 | E | GLN | NE2 | 35 | A | GLU | OE1 | | 2 | 0,136111 | 03:47 | 40:32:00 | 999.99 |
| 500 | E | THR | OG1 | 41 | A | TYR | OH | - | | 02:54 | 0,44375 | 999.99 | 999.99 |

Dd-a = Distance Between Donor and Acceptor
Dh-a = Distance Between Hydrogen and Acceptor
A(d-H-N) = Angle Between Donor-H-N
A(a-O=C) = Angle Between Acceptor-O=C
MO = Multiple Occupancy
Note that angles that are undefined are written as 999.99

**Protein-Protein Ionic Interactions within 6 anstroms**

| Position | Residue | Chain | Position | Residue | Chain |
|---|---|---|---|---|---|
| 30 | ASP | A | 417 | LYS | E |

**Protein-Protein Aromatic-Aromatic Interactions within 4.5 and 7 Angstroms**

| Residue | Position | Chain | Residue | Position | Chain | D(centroid-centroid) | Dihedral Angle |
|---|---|---|---|---|---|---|---|
| 83 | TYR | A | | 486 | PHE | E | 05:00 | 150:00:00 |

**Protein-Protein Aromatic-Sulphur Interactions within 5.3 Angstroms**

| Position | Residue | Chain | Position | Residue | Chain | D(Centroid-Sulphur) | Angle |
|---|---|---|---|---|---|---|---|
| 486 | PHE | E | 82 | MET | A | 0,179861 | 148:44:00 |

**Protein-protein cation-pi interactions**

NO PROTEIN-PROTEIN CATION-PI INTERACTIONS FOUND

# B.1.141
# N439K

**Hydrophobic Interactions within 5 Angstroms**

| Position | Residue | Chain | Position | Residue | Chain |
|---|---|---|---|---|---|
| 28 | PHE | A | 489 | TYR | E |
| 79 | LEU | A | 486 | PHE | E |
| 82 | MET | A | 486 | PHE | E |
| 83 | TYR | A | 486 | PHE | E |

**Protein-Protein disulphide bridges**

**Protein-Protein Main Chain-Main Chain H-Bonds**

| DONOR | | | | ACCEPTOR | | | | PARAMETERS | | | | |
|---|---|---|---|---|---|---|---|---|---|---|---|---|
| POS | CHAIN | RES | ATOM | POS | CHAIN | RES | ATOM | MO | Dd-a | Dh-a | A(d-H-N) | A(a-O=C) |
| 502 | E | GLY | N | 353 | A | LYS | O | - | 2.80 | 1.83 | 170.78 | 154.97 |

Dd-a = Distance Between Donor and Acceptor
Dh-a = Distance Between Hydrogen and Acceptor
A(d-H-N) = Angle Between Donor-H-N
A(a-O=C) = Angle Between Acceptor-O=C
MO = Multiple Occupancy
Note that angles that are undefined are written as 999.99

**Protein-Protein Main Chain-Side Chain Hydrogen Bonds**

| DONOR | | | | ACCEPTOR | | | | PARAMETERS | | | | |
|---|---|---|---|---|---|---|---|---|---|---|---|---|
| POS | CHAIN | RES | ATOM | POS | CHAIN | RES | ATOM | MO | Dd-a | Dh-a | A(d-H-N) | A(a-O=C) |
| 28 | A | PHE | N | 489 | E | TYR | OH | - | 3.48 | 3.80 | 63.94 | 999.99 |
| 41 | A | TYR | OH | 500 | E | THR | O | - | 3.41 | 9.99 | 999.99 | 86.85 |
| 330 | A | ASN | ND2 | 500 | E | THR | O | 1 | 3.48 | 3.00 | 108.43 | 114.52 |
| 330 | A | ASN | ND2 | 500 | E | THR | O | 2 | 3.48 | 3.32 | 90.11 | 114.52 |
| 353 | A | LYS | NZ | 496 | E | GLY | O | - | 2.92 | 9.99 | 999.99 | 120.84 |

| | | | | | | | | | | | | |
|---|---|---|---|---|---|---|---|---|---|---|---|---|
| 355 | A | ASP | OD1 | 500 | E | THR | O | 1 | 3.48 | 3.85 | 62.19 | 162.11 |
| 355 | A | ASP | OD1 | 500 | E | THR | O | 2 | 3.48 | 3.90 | 59.74 | 162.11 |
| 355 | A | ASP | OD2 | 500 | E | THR | O | 1 | 3.36 | 3.75 | 61.20 | 131.30 |
| 355 | A | ASP | OD2 | 500 | E | THR | O | 2 | 3.36 | 3.67 | 64.98 | 131.30 |

Dd-a = Distance Between Donor and Acceptor
Dh-a = Distance Between Hydrogen and Acceptor
A(d-H-N) = Angle Between Donor-H-N
A(a-O=C) = Angle Between Acceptor-O=C
MO = Multiple Occupancy
Note that angles that are undefined are written as 999.99

**Protein-Protein Side Chain-Side Chain Hydrogen Bonds**

| DONOR | | | | ACCEPTOR | | | | PARAMETERS | | | | |
|---|---|---|---|---|---|---|---|---|---|---|---|---|
| POS | CHAIN | RES | ATOM | POS | CHAIN | RES | ATOM | MO | Dd-a | Dh-a | A(d-H-N) | A(a-O=C) |
| 24 | A | GLN | OE1 | 487 | E | ASN | ND2 | 1 | 2.68 | 2.83 | 70.81 | 999.99 |
| 24 | A | GLN | OE1 | 487 | E | ASN | ND2 | 2 | 2.68 | 2.29 | 99.17 | 999.99 |
| 31 | A | LYS | NZ | 493 | E | GLN | OE1 | - | 2.92 | 9.99 | 999.99 | 999.99 |
| 35 | A | GLU | OE2 | 493 | E | GLN | OE1 | 1 | 3.34 | 3.69 | 62.92 | 999.99 |
| 35 | A | GLU | OE2 | 493 | E | GLN | OE1 | 2 | 3.34 | 2.31 | 160.55 | 999.99 |
| 35 | A | GLU | OE2 | 493 | E | GLN | NE2 | 1 | 2.91 | 3.45 | 51.23 | 999.99 |
| 35 | A | GLU | OE2 | 493 | E | GLN | NE2 | 2 | 2.91 | 2.25 | 117.46 | 999.99 |
| 83 | A | TYR | OH | 487 | E | ASN | OD1 | - | 2.74 | 9.99 | 999.99 | 999.99 |
| 417 | E | LYS | NZ | 30 | A | ASP | OD2 | - | 2.58 | 9.99 | 999.99 | 999.99 |
| 449 | E | TYR | OH | 38 | A | ASP | OD1 | - | 2.99 | 9.99 | 999.99 | 999.99 |
| 449 | E | TYR | OH | 38 | A | ASP | OD2 | - | 2.81 | 9.99 | 999.99 | 999.99 |
| 487 | E | ASN | ND2 | 24 | A | GLN | OE1 | 1 | 2.68 | 1.90 | 127.00 | 999.99 |
| 487 | E | ASN | ND2 | 24 | A | GLN | OE1 | 2 | 2.68 | 3.05 | 59.48 | 999.99 |
| 487 | E | ASN | OD1 | 83 | A | TYR | OH | 1 | 2.74 | 2.30 | 102.35 | 999.99 |
| 487 | E | ASN | OD1 | 83 | A | TYR | OH | 2 | 2.74 | 2.60 | 85.67 | 999.99 |
| 493 | E | GLN | OE1 | 35 | A | GLU | OE2 | 1 | 3.34 | 2.52 | 132.95 | 999.99 |
| 493 | E | GLN | OE1 | 35 | A | GLU | OE2 | 2 | 3.34 | 4.27 | 25.99 | 999.99 |

| | | | | | | | | | | | | |
|---|---|---|---|---|---|---|---|---|---|---|---|---|
| 493 | E | GLN | NE2 | 35 | A | GLU | OE2 | 1 | 2.91 | 1.89 | 161.57 | 999.99 |
| 493 | E | GLN | NE2 | 35 | A | GLU | OE2 | 2 | 2.91 | 3.63 | 39.93 | 999.99 |
| 500 | E | THR | OG1 | 41 | A | TYR | OH | - | 2.70 | 9.99 | 999.99 | 999.99 |
| 505 | E | TYR | OH | 37 | A | GLU | OE1 | - | 2.94 | 9.99 | 999.99 | 999.99 |

Dd-a = Distance Between Donor and Acceptor
Dh-a = Distance Between Hydrogen and Acceptor
A(d-H-N) = Angle Between Donor-H-N
A(a-O=C) = Angle Between Acceptor-O=C
MO = Multiple Occupancy
Note that angles that are undefined are written as 999.99

**Protein-Protein Ionic Interactions within 6 anstroms**

| Position | Residue | Chain | Position | Residue | Chain |
|---|---|---|---|---|---|
| 30 | ASP | A | 417 | LYS | E |
| 31 | LYS | A | 484 | GLU | E |

**Protein-Protein Aromatic-Aromatic Interactions within 4.5 and 7 Angstroms**

| Position | Residue | Chain | Position | Residue | Chain | D(centroic | Dihedral Angle |
|---|---|---|---|---|---|---|---|
| 83 | TYR | A | 486 | PHE | E | 5.00 | 148.54 |

**Protein-Protein Aromatic-Sulphur Interactions within 5.3 Angstroms**

| Position | Residue | Chain | Position | Residue | Chain | D(centroic | Angle |
|---|---|---|---|---|---|---|---|
| 486 | PHE | E | 82 | MET | A | 3.83 | 145.26 |

**Protein-protein cation-pi interactions**

#### Summary VoC.spike.RBD/ACE2 interactions

| VoC SARS-CoV-2 spike RBD/ACE2 Interactions | hCoV.19Wuhan.WIV04.2019 (6m0j.pdb) | B.1.1.7_UK S494P_N501Y_E484K | B.1.315_S.africa N501Y_E484K_K417T | B.1.427 California L452R | B1.617.india E484Q_L452R | Japan Brazil N501Y_E484K_K417N | B.1.141 N439K | B.1.620 S477N_E484K |
|---|---|---|---|---|---|---|---|---|
| Interactions within 5 Angstroms | 4 | 5 | 5 | 4 | 4 | 5 | 4 | 4 |
| Protein-Protein disulphide bridges | 0 | 0 | 0 | 0 | 0 | 0 | 0 | 0 |
| Protein-Protein Main Chain-Main Chain H-Bonds | 1 | 1 | 1 | 1 | 1 | 1 | 1 | 1 |
| Protein-Protein Main Chain-Side Chain Hydrogen Bonds | 6 | 2 | 7 | 5 | 3 | 5 | 9 | 4 |
| Protein-Protein Side Chain-Side Chain Hydrogen Bonds | 23 | 14 | 19 | 24 | 18 | 22 | 21 | 23 |
| Protein-Protein Ionic Interactions within 6 anstroms | 2 | 1 | 0 | 3 | 1 | 0 | 2 | 1 |
| Protein-Protein Aromatic-Aromatic Interactions within 4.5 and 7 Angstroms | 1 | 2 | 2 | 1 | 1 | 2 | 1 | 1 |
| Protein-Protein Aromatic-Sulphur Interactions within 5.3 Angstroms | 1 | 1 | 1 | 1 | 1 | 1 | 1 | 1 |
| Protein-protein cation-pi interactions | 0 | 1 | 1 | 0 | 0 | 1 | 0 | 0 |
| Total | 38 | 27 | 36 | 39 | 29 | 37 | 39 | 35 |

### ACE(6h5w.pdb)_Sars-CoV-2.spike.RBD

**Hydrophobic Interactions within 5 Angstroms**

**Protein-Protein disulphide bridges**

**Protein-Protein Main Chain-Main Chain H-Bonds**

**Protein-Protein Main Chain-Side Chain Hydrogen Bond:**

**Protein-Protein Side Chain-Side Chain Hydrogen Bonds**

| DONOR | | | | ACCEPTOR | | | | PARAMETERS | | | | |
|---|---|---|---|---|---|---|---|---|---|---|---|---|
| POS | CHAIN | RES | ATOM | POS | CHAIN | RES | ATOM | MO | Dd-a | Dh-a | A(d-H-N) | A(a-O=C) |
| 43 | A | GLU | OE2 | 487 | E | ASN | ND2 | 1 | 0,147222222 | 03:18 | 66:44:00 | 999.99 |
| 43 | A | GLU | OE2 | 487 | E | ASN | ND2 | 2 | 0,147222222 | 02:27 | 117:28:00 | 999.99 |
| 46 | A | LYS | NZ | 473 | E | TYR | OH | - | 0,145138889 | 0,44375 | 999.99 | 999.99 |
| 53 | A | ARG | NH1 | 493 | E | GLN | OE1 | 1 | 0,146527778 | 02:07 | 134:15:00 | 999.99 |
| 53 | A | ARG | NH1 | 493 | E | GLN | OE1 | 2 | 0,146527778 | 03:39 | 54:00:00 | 999.99 |
| 363 | A | LYS | NZ | 498 | E | GLN | OE1 | - | 03:22 | 0,44375 | 999.99 | 999.99 |
| 417 | E | LYS | NZ | 49 | A | GLU | OE1 | - | 0,129861111 | 0,44375 | 999.99 | 999.99 |
| 487 | E | ASN | ND2 | 43 | A | GLU | OE2 | 1 | 0,147222222 | 0,104861111 | 157.64 | 999.99 |
| 487 | E | ASN | ND2 | 43 | A | GLU | OE2 | 2 | 0,147222222 | 03:46 | 51:39:00 | 999.99 |
| 500 | E | THR | OG1 | 334 | A | GLU | OE1 | - | 03:20 | 0,44375 | 999.99 | 999.99 |
| 500 | E | THR | OG1 | 334 | A | GLU | OE2 | - | 0,142361111 | 0,44375 | 999.99 | 999.99 |
| 505 | E | TYR | OH | 396 | A | ASP | OD1 | - | 0,129166667 | 0,44375 | 999.99 | 999.99 |

Dd-a = Distance Between Donor and Acceptor
Dh-a = Distance Between Hydrogen and Acceptor
A(d-H-N) = Angle Between Donor-H-N
A(a-O=C) = Angle Between Acceptor-O=C
MO = Multiple Occupancy
Note that angles that are undefined are written as 999.99

**Protein-Protein Ionic Interactions within 6 anstroms**

| Position | Residue | Chain | Position | Residue | Chain |
|---|---|---|---|---|---|
| 49 | GLU | A | 417 | LYS | E |
| 53 | ARG | A | 484 | GLU | E |

**Protein-Protein Aromatic-Aromatic Interactions within 4.5 and 7 Angstroms**

**Protein-Protein Aromatic-Sulphur Interactions within 5.3 Angstroms**

**Protein-protein cation-pi interactions**

| Position | Residue | Chain | Position | Residue | Chain | D(cation-Pi) | Angle |
|---|---|---|---|---|---|---|---|
| 473 | TYR | E | 46 | LYS | A | 05:08 | 74:11:00 |
| 486 | PHE | E | 101 | LYS | A | 04:26 | 159.82 |

B.1.620
6m0j.S477N.E484K.post.rep.pdb

**Hydrophobic Interactions within 5 Angstroms**

| Position | Residue | Chain | Position | Residue | Chain |
|---|---|---|---|---|---|

| | | | | | |
|---|---|---|---|---|---|
| 28 | PHE | A | 489 | TYR | E |
| 79 | LEU | A | 486 | PHE | E |
| 82 | MET | A | 486 | PHE | E |
| 83 | TYR | A | 486 | PHE | E |

**Protein-Protein disulphide bridges**

NO PROTEIN-PROTEIN DISULPHIDE BRIDGES FOUND

**Protein-Protein Main Chain-Main Chain Hydrogen Bonds**

| DONOR | | | | ACCEPTOR | | | | PARAMETERS | | | | |
|---|---|---|---|---|---|---|---|---|---|---|---|---|
| POS | CHAIN | RES | ATOM | POS | CHAIN | RES | ATOM | MO | Dd-a | Dh-a | A(d-H-N) | A(a-O=C) |
| 502 | E | GLY | N | 353 | A | LYS | O | - | 0,138888889 | 0,099305556 | 167:12:00 | 153:07:00 |

Dd-a = Distance Between Donor and Acceptor
Dh-a = Distance Between Hydrogen and Acceptor
A(d-H-N) = Angle Between Donor-H-N
A(a-O=C) = Angle Between Acceptor-O=C
MO = Multiple Occupancy
Note that angles that are undefined are written as 999.99

**Protein-Protein Main Chain-Side Chain Hydrogen Bonds**

| DONOR | | | | ACCEPTOR | | | | PARAMETERS | | | | |
|---|---|---|---|---|---|---|---|---|---|---|---|---|
| POS | CHAIN | RES | ATOM | POS | CHAIN | RES | ATOM | MO | Dd-a | Dh-a | A(d-H-N) | A(a-O=C) |
| 41 | A | TYR | OH | 500 | E | THR | O | - | 03:46 | 0,44375 | 999.99 | 74:35:00 |
| 353 | A | LYS | NZ | 496 | E | GLY | O | - | 0,145833333 | 0,44375 | 999.99 | 121:58:00 |
| 355 | A | ASP | OD2 | 500 | E | THR | O | 1 | 03:21 | 0,166666667 | 60:38:00 | 113.88 |
| 355 | A | ASP | OD2 | 500 | E | THR | O | 2 | 03:21 | 03:37 | 72:50:00 | 113.88 |

Dd-a = Distance Between Donor and Acceptor
Dh-a = Distance Between Hydrogen and Acceptor
A(d-H-N) = Angle Between Donor-H-N
A(a-O=C) = Angle Between Acceptor-O=C
MO = Multiple Occupancy
Note that angles that are undefined are written as 999.99

**Protein-Protein Side Chain-Side Chain Hydrogen Bonds**

| DONOR | | | | ACCEPTOR | | | | PARAMETERS | | | | |
|---|---|---|---|---|---|---|---|---|---|---|---|---|
| POS | CHAIN | RES | ATOM | POS | CHAIN | RES | ATOM | MO | Dd-a | Dh-a | A(d-H-N) | A(a-O=C) |
| 24 | A | GLN | OE1 | 487 | E | ASN | ND2 | 1 | 02:58 | 0,132638889 | 71.66 | 999.99 |
| 24 | A | GLN | OE1 | 487 | E | ASN | ND2 | 2 | 02:58 | 02:23 | 96:42:00 | 999.99 |
| 31 | A | LYS | NZ | 493 | E | GLN | OE1 | - | 0,141666667 | 0,44375 | 999.99 | 999.99 |
| 35 | A | GLU | OE1 | 493 | E | GLN | OE1 | 1 | 03:28 | 03:44 | 72:06:00 | 999.99 |
| 35 | A | GLU | OE1 | 493 | E | GLN | OE1 | 2 | 03:28 | 0,166666667 | 64:22:00 | 999.99 |
| 35 | A | GLU | OE1 | 493 | E | GLN | NE2 | 1 | 0,136111111 | 03:09 | 61.99 | 999.99 |
| 35 | A | GLU | OE1 | 493 | E | GLN | NE2 | 2 | 0,136111111 | 02:36 | 100:05:00 | 999.99 |
| 42 | A | GLN | NE2 | 449 | E | TYR | OH | 1 | 03:23 | 03:35 | 74:19:00 | 999.99 |
| 42 | A | GLN | NE2 | 449 | E | TYR | OH | 2 | 03:23 | 02:52 | 123.71 | 999.99 |
| 83 | A | TYR | OH | 487 | E | ASN | OD1 | - | 0,136805556 | 0,44375 | 999.99 | 999.99 |
| 417 | E | LYS | NZ | 30 | A | ASP | OD2 | - | 0,142361111 | 0,44375 | 999.99 | 999.99 |
| 449 | E | TYR | OH | 38 | A | ASP | OD1 | - | 03:05 | 0,44375 | 999.99 | 999.99 |
| 449 | E | TYR | OH | 38 | A | ASP | OD2 | - | 0,128472222 | 0,44375 | 999.99 | 999.99 |
| 449 | E | TYR | OH | 42 | A | GLN | NE2 | - | 03:23 | 0,44375 | 999.99 | 999.99 |
| 487 | E | ASN | ND2 | 24 | A | GLN | OE1 | 1 | 02:58 | 0,092361111 | 133:46:00 | 999.99 |
| 487 | E | ASN | ND2 | 24 | A | GLN | OE1 | 2 | 02:58 | 03:02 | 55.78 | 999.99 |
| 487 | E | ASN | OD1 | 83 | A | TYR | OH | 1 | 0,136805556 | 02:33 | 102:21:00 | 999.99 |
| 487 | E | ASN | OD1 | 83 | A | TYR | OH | 2 | 0,136805556 | 0,135416667 | 79.88 | 999.99 |
| 493 | E | GLN | OE1 | 35 | A | GLU | OE1 | 1 | 03:28 | 02:49 | 129:24:00 | 999.99 |
| 493 | E | GLN | OE1 | 35 | A | GLU | OE1 | 2 | 03:28 | 04:22 | 1,020139 | 999.99 |
| 493 | E | GLN | NE2 | 35 | A | GLU | OE1 | 1 | 0,136111111 | 0,092361111 | 162.71 | 999.99 |

| | | | | | | | | | | | |
|---|---|---|---|---|---|---|---|---|---|---|---|
| 493 | E | GLN | NE2 | 35 | A | GLU | OE1 | 2 | 0,136111111 | 03:47 | 40:32:00 | 999.99 |
| 500 | E | THR | OG1 | 41 | A | TYR | OH | - | 02:54 | 0,44375 | 999.99 | 999.99 |

Dd-a = Distance Between Donor and Acceptor
Dh-a = Distance Between Hydrogen and Acceptor
A(d-H-N) = Angle Between Donor-H-N
A(a-O=C) = Angle Between Acceptor-O=C
MO = Multiple Occupancy
Note that angles that are undefined are written as 999.99

**Protein-Protein Ionic Interactions within 6 anstroms**

| Position | Residue | Chain | Position | Residue | Chain |
|---|---|---|---|---|---|
| 30 | ASP | A | 417 | LYS | E |

**Protein-Protein Aromatic-Aromatic Interactions within 4.5 and 7 Angstroms**

| Residue | Position | Chain | Residue | Position | Chain | D(centroid-centroid) | Dihedral Angle |
|---|---|---|---|---|---|---|---|
| 83 | TYR | A | 486 | PHE | E | 05:00 | 150:00:00 |

**Protein-Protein Aromatic-Sulphur Interactions within 5.3 Angstroms**

| Position | Residue | Chain | Position | Residue | Chain | D(Centroid-Sulphur) | Angle |
|---|---|---|---|---|---|---|---|
| 486 | PHE | E | 82 | MET | A | 0,17986111 | 148:44:00 |

**Protein-protein cation-pi interactions**

NO PROTEIN-PROTEIN CATION-PI INTERACTIONS FOUND

### NLN(1i1i.pdb)_SARS-CoV-2.spike.RBD

**Hydrophobic Interactions within 5 Angstroms**

| Position | Residue | Chain | Position | Residue | Chain |
|---|---|---|---|---|---|
| 14 | MET | P | 508 | TYR | E |
| 24 | VAL | P | 497 | PHE | E |
| 24 | VAL | P | 505 | TYR | E |
| 24 | VAL | P | 507 | PRO | E |
| 48 | VAL | P | 489 | TYR | E |
| 52 | VAL | P | 486 | PHE | E |
| 55 | ILE | P | 486 | PHE | E |
| 57 | LEU | P | 486 | PHE | E |
| 60 | VAL | P | 486 | PHE | E |
| 68 | VAL | P | 456 | PHE | E |
| 68 | VAL | P | 489 | TYR | E |
| 408 | VAL | P | 503 | VAL | E |
| 437 | LEU | P | 505 | TYR | E |
| 438 | PRO | P | 453 | TYR | E |
| 438 | PRO | P | 455 | LEU | E |
| 443 | MET | P | 505 | TYR | E |
| 444 | MET | P | 503 | VAL | E |

**Protein-Protein disulphide bridges**

**Protein-Protein Main Chain-Main Chain H-Bonds**

| DONOR | | | | ACCEPTOR | | | | | PARAMETERS | | | |
|---|---|---|---|---|---|---|---|---|---|---|---|---|
| **POS** | **CHAIN** | **RES** | **ATOM** | **POS** | **CHAIN** | **RES** | **ATOM** | **MO** | **Dd-a** | **Dh-a** | **A(d-H-N)** | **A(a-O=C)** |
| 14 | P | MET | N | 403 | E | ARG | O | - | 0,142361111 | 0,44375 | 999.99 | 999.99 |
| 23 | P | ASN | N | 505 | E | TYR | O | - | 03:31 | 03:27 | 84:03:00 | 102:45:00 |
| 52 | P | VAL | N | 485 | E | GLY | O | - | 03:26 | 03:47 | 69:58:00 | 123.98 |

| | | | | | | | | | | | | |
|---|---|---|---|---|---|---|---|---|---|---|---|---|
| 53 | P | GLY | N | 485 | E | GLY | O | - | 0,147222222 | 0,110416667 | 158:31:00 | 143:48:00 |
| 54 | P | THR | N | 485 | E | GLY | O | - | 0,148611111 | 0,109722222 | 168:56:00 | 110.87 |
| 406 | E | GLU | N | 14 | P | MET | O | - | 03:34 | 03:42 | 77:12:00 | 163:49:00 |
| 407 | E | VAL | N | 14 | P | MET | O | - | 0,150694444 | 02:05 | 158:56:00 | 144:52:00 |
| 417 | E | LYS | N | 439 | P | ASP | O | - | 03:02 | 02:58 | 107:57:00 | 116.75 |
| 418 | E | ILE | N | 439 | P | ASP | O | - | 0,15 | 02:15 | 141:03:00 | 141.79 |
| 477 | E | SER | N | 56 | P | ALA | O | - | 03:39 | 02:47 | 155:23:00 | 145:23:00 |
| 485 | E | GLY | N | 50 | P | ASP | O | - | 03:40 | 0,168055556 | 69:51:00 | 129:40:00 |
| 486 | E | PHE | N | 51 | P | THR | O | - | 03:46 | 04:10 | 43.79 | 102:51:00 |
| 487 | E | ASN | N | 51 | P | THR | O | - | 03:08 | 02:52 | 115:39:00 | 148:21:00 |
| 488 | E | CYS | N | 51 | P | THR | O | - | 03:50 | 02:54 | 157.99 | 163.61 |
| 500 | E | THR | N | 23 | P | ASN | O | - | 03:06 | 03:17 | 74.63 | 109.86 |
| 500 | E | THR | N | 25 | P | LEU | O | - | 0,147222222 | 02:20 | 130:52:00 | 144:36:00 |
| 503 | E | VAL | N | 444 | P | MET | O | - | 03:05 | 02:10 | 166:04:00 | 156:20:00 |
| 505 | E | TYR | N | 22 | P | ARG | O | - | 03:08 | 02:35 | 131:55:00 | 136.61 |
| 506 | E | GLN | N | 22 | P | ARG | O | - | 03:45 | 0,175 | 66:55:00 | 80.99 |

Dd-a = Distance Between Donor and Acceptor
Dh-a = Distance Between Hydrogen and Acceptor
A(d-H-N) = Angle Between Donor-H-N
A(a-O=C) = Angle Between Acceptor-O=C
MO = Multiple Occupancy
Note that angles that are undefined are written as 999.99

**Protein-Protein Main Chain-Side Chain Hydrogen Bonds**

| DONOR | | | | ACCEPTOR | | | | | PARAMETERS | | | |
|---|---|---|---|---|---|---|---|---|---|---|---|---|
| POS | CHAIN | RES | ATOM | POS | CHAIN | RES | ATOM | MO | Dd-a | Dh-a | A(d-H-N) | A(a-O=C) |
| 20 | P | ALA | N | 506 | E | GLN | NE2 | - | 03:37 | 03:01 | 102:56:00 | 149:36:00 |
| 22 | P | ARG | NH2 | 404 | E | GLY | O | 1 | 0,140277778 | 03:25 | 56:58:00 | 133:32:00 |
| 22 | P | ARG | NH2 | 404 | E | GLY | O | 2 | 0,140277778 | 0,104166667 | 143.93 | 133:32:00 |
| 22 | P | ARG | NE | 503 | E | VAL | O | - | 0,136111111 | 0,104166667 | 138:22:00 | 157:24:00 |
| 22 | P | ARG | NH2 | 503 | E | VAL | O | 1 | 0,136111111 | 0,102777778 | 139.92 | 153:19:00 |

| | | | | | | | | | | | | | |
|---|---|---|---|---|---|---|---|---|---|---|---|---|---|
| 22 | P | ARG | NH2 | 503 | E | VAL | O | | 2 | 0,136111111 | 0,169444444 | 28:59:00 | 153:19:00 |
| 23 | P | ASN | OD1 | 499 | E | PRO | O | | 1 | 03:10 | 0,146527778 | 90:30:00 | 124:01:00 |
| 23 | P | ASN | OD1 | 499 | E | PRO | O | | 2 | 03:10 | 03:55 | 57:15:00 | 124:01:00 |
| 23 | P | ASN | ND2 | 500 | E | THR | O | | 1 | 03:30 | 0,190972222 | 45.92 | 87.70 |
| 23 | P | ASN | ND2 | 500 | E | THR | O | | 2 | 03:30 | 0,125694444 | 122.93 | 87.70 |
| 51 | P | THR | OG1 | 488 | E | CYS | O | - | | 0,139583333 | 0,44375 | 999.99 | 104.82 |
| 54 | P | THR | OG1 | 485 | E | GLY | O | - | | 03:43 | 0,44375 | 999.99 | 68:32:00 |
| 58 | P | LYS | N | 477 | E | SER | OG | - | | 03:13 | 0,145833333 | 94.76 | 999.99 |
| 59 | P | GLU | N | 477 | E | SER | OG | - | | 03:00 | 02:03 | 167.99 | 999.99 |
| 61 | P | THR | N | 487 | E | ASN | OD1 | - | | 03:17 | 02:24 | 154:20:00 | 135:45:00 |
| 64 | P | ASN | OD1 | 487 | E | ASN | O | | 1 | 03:27 | 03:41 | 73:26:00 | 69.99 |
| 64 | P | ASN | OD1 | 487 | E | ASN | O | | 2 | 03:27 | 02:37 | 140:55:00 | 69.99 |
| 413 | P | TYR | OH | 500 | E | THR | O | - | | 03:23 | 0,44375 | 999.99 | 135.76 |
| 432 | P | GLN | OE1 | 502 | E | GLY | O | | 1 | 03:15 | 02:39 | 127:34:00 | 97:06:00 |
| 432 | P | GLN | OE1 | 502 | E | GLY | O | | 2 | 03:15 | 0,193055556 | 34:08:00 | 97:06:00 |
| 432 | P | GLN | NE2 | 502 | E | GLY | O | | 1 | 03:03 | 02:22 | 132:59:00 | 116:47:00 |
| 432 | P | GLN | NE2 | 502 | E | GLY | O | | 2 | 03:03 | 0,181944444 | 36:45:00 | 116:47:00 |
| 404 | E | GLY | N | 14 | P | MET | SD | - | | 0,185416667 | 0,220138889 | 22:16 | 999.99 |
| 404 | E | GLY | N | 441 | P | SER | OG | - | | 03:31 | 02:52 | 136.61 | 999.99 |
| 409 | E | GLN | NE2 | 440 | P | GLY | O | | 1 | 03:37 | 0,147222222 | 106:48:00 | 92:29:00 |
| 409 | E | GLN | NE2 | 440 | P | GLY | O | | 2 | 03:37 | 03:11 | 95:37:00 | 92:29:00 |
| 417 | E | LYS | NZ | 438 | P | PRO | O | - | | 0,148611111 | 0,44375 | 999.99 | 150:16:00 |
| 437 | E | ASN | OD1 | 20 | P | ALA | O | | 1 | 03:04 | 0,141666667 | 90:07:00 | 165.91 |
| 437 | E | ASN | OD1 | 20 | P | ALA | O | | 2 | 03:04 | 03:57 | 52:18:00 | 165.91 |
| 437 | E | ASN | ND2 | 20 | P | ALA | O | | 1 | 03:32 | 03:16 | 89.61 | 125:57:00 |
| 437 | E | ASN | ND2 | 20 | P | ALA | O | | 2 | 03:32 | 0,19375 | 44:27:00 | 125:57:00 |
| 477 | E | SER | OG | 56 | P | ALA | O | - | | 02:56 | 0,44375 | 999.99 | 106.95 |
| 485 | E | GLY | N | 51 | P | THR | OG1 | - | | 03:07 | 02:17 | 151.91 | 999.99 |
| 487 | E | ASN | ND2 | 64 | P | ASN | O | | 1 | 03:10 | 03:44 | 62.78 | 92.84 |
| 487 | E | ASN | ND2 | 64 | P | ASN | O | | 2 | 03:10 | 02:38 | 123.95 | 92.84 |
| 488 | E | CYS | N | 64 | P | ASN | OD1 | - | | 03:19 | 03:56 | 60:32:00 | 166:50:00 |
| 498 | E | GLN | NE2 | 24 | P | VAL | O | | 1 | 0,14375 | 03:40 | 52:07:00 | 124.88 |
| 498 | E | GLN | NE2 | 24 | P | VAL | O | | 2 | 0,14375 | 0,1 | 167.75 | 124.88 |
| 500 | E | THR | OG1 | 389 | P | HIS | O | - | | 0,140277778 | 0,44375 | 999.99 | 112.97 |
| 504 | E | GLY | N | 432 | P | GLN | OE1 | - | | 03:48 | 02:57 | 155:07:00 | 149:50:00 |

| | | | | | | | | | | | | |
|---|---|---|---|---|---|---|---|---|---|---|---|---|
| 506 | E | GLN | N | 14 | P | MET | SD | - | 03:28 | 02:41 | 149.82 | 999.99 |
| 506 | E | GLN | NE2 | 20 | P | ALA | O | 1 | 0,136111111 | 03:44 | 42.98 | 113:05:00 |
| 506 | E | GLN | NE2 | 20 | P | ALA | O | 2 | 0,136111111 | 0,091666667 | 165:01:00 | 113:05:00 |

Dd-a = Distance Between Donor and Acceptor
Dh-a = Distance Between Hydrogen and Acceptor
A(d-H-N) = Angle Between Donor-H-N
A(a-O=C) = Angle Between Acceptor-O=C
MO = Multiple Occupancy
Note that angles that are undefined are written as 999.99

**Protein-Protein Side Chain-Side Chain Hydrogen Bonds**

| DONOR | | | | ACCEPTOR | | | | | PARAMETERS | | | |
|---|---|---|---|---|---|---|---|---|---|---|---|---|
| **POS** | **CHAIN** | **RES** | **ATOM** | **POS** | **CHAIN** | **RES** | **ATOM** | **MO** | **Dd-a** | **Dh-a** | **A(d-H-N)** | **A(a-O=C)** |
| 16 | P | SER | OG | 405 | E | ASP | OD1 | - | 0,147916667 | 0,44375 | 999.99 | 999.99 |
| 18 | P | THR | OG1 | 506 | E | GLN | OE1 | - | 0,134027778 | 0,44375 | 999.99 | 999.99 |
| 37 | P | ARG | NH1 | 449 | E | TYR | OH | 1 | 03:36 | 03:32 | 82.69 | 999.99 |
| 37 | P | ARG | NH1 | 449 | E | TYR | OH | 2 | 03:36 | 0,177777778 | 58.86 | 999.99 |
| 51 | P | THR | OG1 | 484 | E | GLU | OE1 | - | 02:49 | 0,44375 | 999.99 | 999.99 |
| 432 | P | GLN | NE2 | 501 | E | ASN | OD1 | 1 | 02:59 | 03:11 | 52:03:00 | 999.99 |
| 432 | P | GLN | NE2 | 501 | E | ASN | OD1 | 2 | 02:59 | 0,086805556 | 146.84 | 999.99 |
| 441 | P | SER | OG | 406 | E | GLU | OE1 | - | 03:49 | 0,44375 | 999.99 | 999.99 |
| 403 | E | ARG | NH1 | 439 | P | ASP | OD2 | 1 | 0,143055556 | 0,098611111 | 165:42:00 | 999.99 |
| 403 | E | ARG | NH1 | 439 | P | ASP | OD2 | 2 | 0,143055556 | 03:39 | 50.98 | 999.99 |
| 417 | E | LYS | NZ | 67 | P | GLN | OE1 | - | 03:00 | 0,44375 | 999.99 | 999.99 |
| 500 | E | THR | OG1 | 413 | P | TYR | OH | - | 0,134027778 | 0,44375 | 999.99 | 999.99 |
| 501 | E | ASN | OD1 | 432 | P | GLN | NE2 | 1 | 02:59 | 0,147916667 | 61:26:00 | 999.99 |
| 501 | E | ASN | OD1 | 432 | P | GLN | NE2 | 2 | 02:59 | 02:25 | 96:29:00 | 999.99 |
| 506 | E | GLN | OE1 | 18 | P | THR | OG1 | 1 | 0,134027778 | 02:00 | 122:39:00 | 999.99 |
| 506 | E | GLN | OE1 | 18 | P | THR | OG1 | 2 | 0,134027778 | 03:08 | 61:17:00 | 999.99 |

Dd-a = Distance Between Donor and Acceptor
Dh-a = Distance Between Hydrogen and Acceptor
A(d-H-N) = Angle Between Donor-H-N
A(a-O=C) = Angle Between Acceptor-O=C
MO = Multiple Occupancy
Note that angles that are undefined are written as 999.99

**Protein-Protein Ionic Interactions within 6 anstroms**

| Position | Residue | Chain | Position | Residue | Chain |
|---|---|---|---|---|---|
| 71 | ASP | P | 417 | LYS | E |
| 439 | ASP | P | 403 | ARG | E |

**Protein-Protein Aromatic-Aromatic Interactions within 4.5 and 7 Angstroms**

**Protein-Protein Aromatic-Sulphur Interactions within 5.3 Angstroms**

**Protein-protein cation-pi interactions**

| Position | Residue | Chain | Position | Residue | Chain | D(cation-Pi) | Angle |
|---|---|---|---|---|---|---|---|
| 449 | TYR | E | 37 | ARG | P | 0,19375 | 144.84 |

### THOP(1s4b.pdb)_Sars-CoV-2.spike.RBD

**Hydrophobic Interactions within 5 Angstroms**

| Position | Residue | Chain | Position | Residue | Chain |
|---|---|---|---|---|---|
| 28 | LEU | P | 445 | VAL | E |
| 67 | ALA | P | 455 | LEU | E |
| 67 | ALA | P | 489 | TYR | E |
| 73 | VAL | P | 505 | TYR | E |
| 77 | VAL | P | 505 | TYR | E |
| 124 | TRP | P | 486 | PHE | E |
| 407 | VAL | P | 503 | VAL | E |
| 443 | ILE | P | 503 | VAL | E |

**Protein-Protein disulphide bridges**

**Protein-Protein Main Chain-Main Chain H-Bonds**

| DONOR | | | | ACCEPTOR | | | | | PARAMETERS | | | |
|---|---|---|---|---|---|---|---|---|---|---|---|---|
| **POS** | **CHAIN** | **RES** | **ATOM** | **POS** | **CHAIN** | **RES** | **ATOM** | **MO** | **Dd-a** | **Dh-a** | **A(d-H-N)** | **A(a-O=C)** |
| 24 | P | LEU | N | 500 | E | THR | O | - | 0,145138889 | 0,44375 | 999.99 | 999.99 |
| 57 | P | GLU | N | 478 | E | THR | O | - | 0,149305556 | 0,131944444 | 95:22:00 | 131:11:00 |
| 417 | E | LYS | N | 438 | P | ASP | O | - | 03:35 | 03:03 | 100:58:00 | 115.65 |
| 418 | E | ILE | N | 438 | P | ASP | O | - | 0,140277778 | 02:05 | 135:11:00 | 124.87 |
| 475 | E | ALA | N | 57 | P | GLU | O | - | 0,145833333 | 0,105555556 | 164:41:00 | 140:14:00 |
| 476 | E | GLY | N | 57 | P | GLU | O | - | 0,132638889 | 0,095833333 | 152:48:00 | 112:18:00 |
| 477 | E | SER | N | 56 | P | PHE | O | - | 0,148611111 | 02:03 | 154:13:00 | 126:36:00 |
| 485 | E | GLY | N | 50 | P | GLN | O | - | 03:04 | 0,126388889 | 106:09:00 | 108.94 |
| 500 | E | THR | N | 24 | P | LEU | O | - | 0,151388889 | 02:03 | 166.65 | 133:14:00 |
| 502 | E | GLY | N | 443 | P | ILE | O | - | 03:37 | 0,140972222 | 116:27:00 | 139:18:00 |

Dd-a = Distance Between Donor and Acceptor
Dh-a = Distance Between Hydrogen and Acceptor

A(d-H-N) = Angle Between Donor-H-N
A(a-O=C) = Angle Between Acceptor-O=C
MO = Multiple Occupancy
Note that angles that are undefined are written as 999.99

**Protein-Protein Main Chain-Side Chain Hydrogen Bonds**

| DONOR | | | | ACCEPTOR | | | | | PARAMETERS | | | |
|---|---|---|---|---|---|---|---|---|---|---|---|---|
| POS | CHAIN | RES | ATOM | POS | CHAIN | RES | ATOM | MO | Dd-a | Dh-a | A(d-H-N) | A(a-O=C) |
| 24 | P | LEU | N | 501 | E | ASN | ND2 | - | 03:31 | 0,44375 | 999.99 | 135.91 |
| 54 | P | GLN | OE1 | 478 | E | THR | O | 1 | 03:45 | 03:16 | 96.62 | 163:24:00 |
| 54 | P | GLN | OE1 | 478 | E | THR | O | 2 | 03:45 | 03:03 | 104:37:00 | 163:24:00 |
| 54 | P | GLN | NE2 | 486 | E | PHE | O | 1 | 03:50 | 0,168055556 | 75:10:00 | 124:44:00 |
| 54 | P | GLN | NE2 | 486 | E | PHE | O | 2 | 03:50 | 0,174305556 | 70:27:00 | 124:44:00 |
| 57 | P | GLU | OE2 | 476 | E | GLY | O | 1 | 03:39 | 03:24 | 88.69 | 86.60 |
| 57 | P | GLU | OE2 | 476 | E | GLY | O | 2 | 03:39 | 0,128472222 | 125:43:00 | 86.60 |
| 58 | P | ASP | OD2 | 473 | E | TYR | O | 1 | 03:44 | 04:29 | 32.92 | 80:41:00 |
| 58 | P | ASP | OD2 | 473 | E | TYR | O | 2 | 03:44 | 02:51 | 144.95 | 80:41:00 |
| 128 | P | LYS | NZ | 475 | E | ALA | O | - | 0,14375 | 0,44375 | 999.99 | 146:53:00 |
| 441 | P | ARG | N | 405 | E | ASP | OD2 | - | 0,138888889 | 02:08 | 127:55:00 | 128:05:00 |
| 405 | E | ASP | OD2 | 441 | P | ARG | O | 1 | 03:49 | 04:26 | 38:06:00 | 84:35:00 |
| 405 | E | ASP | OD2 | 441 | P | ARG | O | 2 | 03:49 | 03:15 | 99:48:00 | 84:35:00 |
| 417 | E | LYS | NZ | 436 | P | ARG | O | - | 0,145833333 | 0,44375 | 999.99 | 123:36:00 |
| 417 | E | LYS | NZ | 437 | P | GLN | O | - | 03:20 | 0,44375 | 999.99 | 88:05:00 |
| 474 | E | GLN | N | 58 | P | ASP | OD2 | - | 03:38 | 0,191666667 | 47.77 | 133:28:00 |
| 477 | E | SER | N | 57 | P | GLU | OE2 | - | 03:40 | 0,184722222 | 55:39:00 | 138.69 |
| 478 | E | THR | N | 57 | P | GLU | OE2 | - | 03:45 | 03:45 | 81.86 | 93.92 |
| 500 | E | THR | OG1 | 24 | P | LEU | O | - | 03:40 | 0,44375 | 999.99 | 91:55:00 |

Dd-a = Distance Between Donor and Acceptor
Dh-a = Distance Between Hydrogen and Acceptor
A(d-H-N) = Angle Between Donor-H-N

A(a-O=C) = Angle Between Acceptor-O=C
MO = Multiple Occupancy
Note that angles that are undefined are written as 999.99

**Protein-Protein Side Chain-Side Chain Hydrogen Bonds**

| DONOR | | | | ACCEPTOR | | | | | PARAMETERS | | | |
|---|---|---|---|---|---|---|---|---|---|---|---|---|
| POS | CHAIN | RES | ATOM | POS | CHAIN | RES | ATOM | MO | Dd-a | Dh-a | A(d-H-N) | A(a-O=C) |
| 26 | P | TRP | NE1 | 500 | E | THR | OG1 | - | 03:27 | 0,150694444 | 101:03:00 | 999.99 |
| 46 | P | ARG | NE | 484 | E | GLU | OE2 | - | 0,142361111 | 02:10 | 128:55:00 | 999.99 |
| 50 | P | GLN | NE2 | 484 | E | GLU | OE2 | 1 | 0,147222222 | 03:33 | 58:59:00 | 999.99 |
| 50 | P | GLN | NE2 | 484 | E | GLU | OE2 | 2 | 0,147222222 | 0,104861111 | 161:01:00 | 999.99 |
| 54 | P | GLN | NE2 | 480 | E | CYS | SG | 1 | 03:13 | 03:50 | 61:11:00 | 999.99 |
| 54 | P | GLN | NE2 | 480 | E | CYS | SG | 2 | 03:13 | 0,125694444 | 109.98 | 999.99 |
| 54 | P | GLN | NE2 | 488 | E | CYS | SG | 1 | 0,168055556 | 03:52 | 86:45:00 | 999.99 |
| 54 | P | GLN | NE2 | 488 | E | CYS | SG | 2 | 0,168055556 | 03:02 | 116.98 | 999.99 |
| 64 | P | THR | OG1 | 487 | E | ASN | OD1 | - | 02:53 | 0,44375 | 999.99 | 999.99 |
| 78 | P | GLN | OE1 | 449 | E | TYR | OH | 1 | 03:48 | 0,167361111 | 74.63 | 999.99 |
| 78 | P | GLN | OE1 | 449 | E | TYR | OH | 2 | 03:48 | 0,126388889 | 137:42:00 | 999.99 |
| 436 | P | ARG | NH2 | 453 | E | TYR | OH | 1 | 03:24 | 03:08 | 89:37:00 | 999.99 |
| 436 | P | ARG | NH2 | 453 | E | TYR | OH | 2 | 03:24 | 0,16875 | 60:24:00 | 999.99 |
| 437 | P | GLN | OE1 | 453 | E | TYR | OH | 1 | 03:02 | 0,135416667 | 93.76 | 999.99 |
| 437 | P | GLN | OE1 | 453 | E | TYR | OH | 2 | 03:02 | 03:24 | 68:50:00 | 999.99 |
| 437 | P | GLN | NE2 | 493 | E | GLN | OE1 | 1 | 0,144444444 | 03:33 | 56:06:00 | 999.99 |
| 437 | P | GLN | NE2 | 493 | E | GLN | OE1 | 2 | 0,144444444 | 0,100694444 | 165:45:00 | 999.99 |
| 417 | E | LYS | NZ | 70 | P | ASP | OD2 | - | 03:34 | 0,44375 | 999.99 | 999.99 |
| 449 | E | TYR | OH | 39 | P | GLU | OE1 | - | 03:46 | 0,44375 | 999.99 | 999.99 |
| 449 | E | TYR | OH | 78 | P | GLN | OE1 | - | 03:48 | 0,44375 | 999.99 | 999.99 |
| 453 | E | TYR | OH | 437 | P | GLN | OE1 | - | 03:02 | 0,44375 | 999.99 | 999.99 |
| 480 | E | CYS | SG | 54 | P | GLN | NE2 | - | 03:13 | 0,44375 | 999.99 | 999.99 |
| 480 | E | CYS | SG | 58 | P | ASP | OD1 | - | 0,186805556 | 0,44375 | 999.99 | 999.99 |
| 484 | E | GLU | OE2 | 50 | P | GLN | NE2 | 1 | 0,147222222 | 02:19 | 124:48:00 | 999.99 |
| 484 | E | GLU | OE2 | 50 | P | GLN | NE2 | 2 | 0,147222222 | 03:23 | 64:11:00 | 999.99 |

| | | | | | | | | | | | | | |
|---|---|---|---|---|---|---|---|---|---|---|---|---|---|
| 487 | E | ASN | OD1 | 64 | P | THR | OG1 | | 1 | 02:53 | 0,130555556 | 70:24:00 | 999.99 |
| 487 | E | ASN | OD1 | 64 | P | THR | OG1 | | 2 | 02:53 | 0,107638889 | 110:20:00 | 999.99 |
| 488 | E | CYS | SG | 54 | P | GLN | NE2 | - | | 0,168055556 | 0,44375 | 999.99 | 999.99 |
| 488 | E | CYS | SG | 58 | P | ASP | OD1 | - | | 03:20 | 0,44375 | 999.99 | 999.99 |
| 488 | E | CYS | SG | 58 | P | ASP | OD2 | - | | 03:26 | 0,44375 | 999.99 | 999.99 |
| 493 | E | GLN | OE1 | 437 | P | GLN | NE2 | | 1 | 0,144444444 | 03:23 | 61.63 | 999.99 |
| 493 | E | GLN | OE1 | 437 | P | GLN | NE2 | | 2 | 0,144444444 | 02:29 | 112.72 | 999.99 |

Dd-a = Distance Between Donor and Acceptor
Dh-a = Distance Between Hydrogen and Acceptor
A(d-H-N) = Angle Between Donor-H-N
A(a-O=C) = Angle Between Acceptor-O=C
MO = Multiple Occupancy
Note that angles that are undefined are written as 999.99

**Protein-Protein Ionic Interactions within 6 anstroms**

| Position | Residue | Chain | Position | Residue | Chain |
|---|---|---|---|---|---|
| 39 | GLU | P | 444 | LYS | E |
| 46 | ARG | P | 484 | GLU | E |
| 70 | ASP | P | 417 | LYS | E |
| 406 | GLU | P | 408 | ARG | E |
| 438 | ASP | P | 403 | ARG | E |

**Protein-Protein Aromatic-Aromatic Interactions within 4.5 and 7 Angstroms**

**Protein-Protein Aromatic-Sulphur Interactions within 5.3 Angstroms**

**Protein-protein cation-pi interactions**

| Position | Residue | Chain | Position | Residue | Chain | D(cation-Pi) | Angle |
|---|---|---|---|---|---|---|---|
| 453 | TYR | E | 436 | ARG | P | 0,27014 | 111:48:00 |
| 505 | TYR | E | 436 | ARG | P | 0,2625 | 66:51:00 |

**Summary SARS-CoV-2 spike RBD/ACE2 structurally related receptors interactions**

| | SARS-CoV-2 spike RBD/ACE2 (6m0j.pdb) | SARS-CoV-2 spike RBD/ACE (6h5w) | SARS-CoV-2 spike RBD/NLN (1i1i) | SARS-CoV-2 spike RBD/THOP1 (1s4b) |
|---|---|---|---|---|
| **Hydrophobic Interactions within 5 Angstroms** | 4 | 0 | 17 | 8 |
| **Protein-Protein disulphide bridges** | 0 | 0 | 0 | 0 |
| **Protein-Protein Main Chain-Main Chain H-Bonds** | 1 | 0 | 19 | 10 |
| **Protein-Protein Main Chain-Side Chain Hydrogen Bonds** | 6 | 0 | 43 | 19 |
| **Protein-Protein Side Chain-Side Chain Hydrogen Bonds** | 23 | 12 | 16 | 32 |
| **Protein-Protein Ionic Interactions within 6 anstroms** | 2 | 1 | 2 | 5 |
| **Protein-Protein Aromatic-Aromatic Interactions within 4.5 and 7 Angstroms** | 1 | 0 | 0 | 0 |

| | | | | |
|---|---|---|---|---|
| **Protein-Protein Aromatic-Sulphur Interactions within 5.3 Angstroms** | 1 | 0 | 0 | 0 |
| **Protein-protein cation-pi interactions** | 0 | 2 | 1 | 2 |
