## Supplementary material for "A modular molecular framework for quickly estimating the binding affinity of the spike protein of SARS-CoV-2 variants for ACE2, in presence of mutations at the spike receptor binding domain": Supp. Tab. 2

**Supp. Tab. 2 ACE2 structurally related proteins sampled by pGenThreader and I-TASSER**

| **PDB_ID** | **Chain type** | **Chains** | **Target Length** | **protein name** | **virus strain/infected organism** | **RMSD ACE2 (6m0j)** | **RMSD ACE1 (6h5w)** | **RMSD THOP1 (1s4b)** | **RMSD NLN (1i1i)** |
|---|---|---|---|---|---|---|---|---|---|
| 6m0j | Angiotensin-converting enzyme 2 | 2 | 597 | Structure of SARS-CoV-2 chimeric receptor-binding domain complexed with its receptor human ACE2 | *Homo sapiens* | 0 | 2,384 | 5,003 | 3,871 |
| 6zpq | Angiotensin-converting enzyme | 4 | 629 | Crystal structure of the open conformation of Angiotensin-1 converting enzyme N-domain. | *Homo sapiens* | 1,001 | 1,298 | 8,74 | 4,258 |
| 1r42 | angiotensin I converting enzyme 2 | 1 | 615 | Native Human Angiotensin Converting Enzyme-Related Carboxypeptidase (ACE2) | *Homo sapiens* | 0,48 | 3,051 | 5,125 | 4,329 |
| 1r4l | angiotensin I converting enzyme 2 | 1 | 615 | Inhibitor Bound Human Angiotensin Converting Enzyme-Related Carboxypeptidase (ACE2) | *Homo sapiens* | 1,613 | 1,135 | 5,307 | 5,016 |
| 2c6f | ANGIOTENSIN-CONVERTING ENZYME, SOMATIC ISOFORM | 2 | 612 | Structure of human somatic angiontensin-I converting enzyme N domain | *Homo sapiens* | 1,949 | 0,632 | 5,145 | 4,365 |

| | | | | | | | | | |
|---|---|---|---|---|---|---|---|---|---|
| 3nxq | Angiotensin-converting enzyme | 2 | 629 | Angiotensin Converting Enzyme N domain glycsoylation mutant (Ndom389) in complex with RXP407 | *Homo sapiens* | 2,218 | 0,594 | 4,611 | 4,361 |
| 1uzf | ANGIOTENSIN CONVERTING ENZYME | 1 | 589 | Complex of the anti-hypertensive drug captopril an the human testicular angiotensin I-converting enzyme | *Homo sapiens* | 2,233 | 0,236 | 5,434 | 4,451 |
| 5am9 | ANGIOTENSIN-CONVERTING ENZYME | 4 | 629 | Crystal structure of the Angiotensin-1 converting enzyme N-domain in complex with amyloid-beta 10-16 | *Homo sapiens* | 2,282 | 0,627 | 4,705 | 4,648 |
| 5amb | ANGIOTENSIN-CONVERTING ENZYME, AMYLOID BETA A4 PROTEIN | 2, 2 | 629, 8 | Crystal structure of the Angiotensin-1 converting enzyme N-domain in complex with amyloid-beta 35-42 | *Homo sapiens* | 2,293 | 0,561 | 5,317 | 4,979 |
| 3BKLA | Angiotensin-converting enzyme, somatic isoform | 1 | 591 | Testis ACE co-crystal structure with ketone ACE inhibitor kAW | *Homo sapiens* | 2,337 | 0,231 | 5,371 | 4,497 |

| | | | | | | | | | |
|---|---|---|---|---|---|---|---|---|---|
| 4bzr | ANGIOTENSIN-CONVERTING ENZYME | 1 | 589 | Human testis angiotensin converting enzyme in complex with K-26 | *Homo sapiens* | 2,357 | 0,19 | 5,168 | 4,039 |
| 3bkk | Angiotensin-converting enzyme, somatic isoform | 1 | 591 | Tesis ACE co-crystal structure with ketone ACE inhibitor kAF | *Homo sapiens* | 2,367 | 0,226 | 5,494 | 5,046 |
| 6h5w | Angiotensin-converting enzyme | 1 | 591 | Crystal structure of human Angiotensin-1 converting enzyme C-domain in complex with Omapatrilat. | *Homo sapiens* | 2,83 | 0 | 5,5 | 5,191 |
| 1j36 | angiotensin converting enzyme | 2 | 607 | Crystal Structure of Drosophila AnCE | *Drosophila melanogaster* | 2,388 | 0,792 | 4,885 | 4,33 |
| 1o86A | ANGIOTENSIN CONVERTING ENZYME | 1 | 589 | Crystal Structure of Human Angiotensin Converting Enzyme in complex with lisinopril. | *Homo sapiens* | 2,446 | 0,214 | 5,641 | 4,235 |
| 4ca7 | ANGIOTENSIN-CONVERTING ENZYME | 1 | 598 | Drosophila Angiotensin converting enzyme (AnCE) in complex with a phosphinic tripeptide FI | *Drosophila melanogaster* | 2,463 | 0,833 | 5,148 | 4,327 |

| | | | | | | | | | |
|---|---|---|---|---|---|---|---|---|---|
| 2xy9A | ANGIOTENSIN-CONVERTING ENZYME | 1 | 585 | Human Angiotensin converting enzyme in complex with phosphinic tripeptide | *Homo sapiens* | 2,494 | 0,336 | 5,164 | 4,339 |
| 6s1y | Angiotensin-converting enzyme | 1 | 621 | Crystal structure of Anopheles gambiae AnoACE2 in complex with gamma-Polyglutamic Acid. | *Anopheles gambiae* | 2,511 | 0,929 | 5,608 | 4,819 |
| 2o3e | Neurolysin | 1 | 678 | Crystal structure of engineered neurolysin with thimet oligopeptidase specificity for neurotensin cleavage site. | *Rattus Norvegicus* | 3,84 | 5,172 | 1,087 | 0,216 |
| 1i1i | NEUROLYSIN | 1 | 681 | NEUROLYSIN (ENDOPEPTIDASE 24.16) CRYSTAL STRUCTURE | *Rattus norvegicus* | 3,9 | 5,191 | 1,116 | 0 |
| 5wvu | Thermostable carboxypeptidase 1 | 3 | 510 | Crystal structure of carboxypeptidase from Thermus thermophilus | *Thermus thermophilus HB8* | 4,678 | 6,643 | 5,686 | 4,35 |

| | | | | | | | | | |
|---|---|---|---|---|---|---|---|---|---|
| 3hoa | Thermostable carboxypeptidase 1 | 2 | 509 | Crystal structure of the Thermus thermophilus M32 carboxypeptidase | *Thermus thermophilus HB27* | 4,909 | 6,272 | 5,999 | 5,028 |
| 1ka2 | M32 carboxypeptidase | 1 | 499 | Structure of Pyrococcus furiosus Carboxypeptidase Apo-Mg | *Pyrococcus furiosus* | 4,988 | 5.641 | 4,754 | 4,585 |
| 3ce2 | Putative peptidase | 1 | 618 | Crystal structure of putative peptidase from Chlamydophila abortus | *Chlamydia abortus S26/3* | 4,991 | 8,426 | 6,023 | 6,407 |
| 1s4b | Thimet oligopeptidase | 1 | 674 | Crystal structure of human thimet oligopeptidase. | *Homo sapiens* | 4,46 | 5,5 | 0 | 1,116 |
| 3dwc | Metallocarboxypeptidase | 3 | 505 | Trypanosoma Cruzi Metallocarboxypeptidase 1 | *Trypanosoma cruzi* | 5,082 | 4,872 | 5,987 | 5,584 |
| 5e3x | Thermostable carboxypeptidase 1 | 1 | 489 | Crystal structure of thermostable Carboxypeptidase (FisCP) from Fervidobacterium Islandicum AW-1 | *Fervidobacterium islandicum* | 5,242 | 4,179 | 5,634 | 5,778 |
| 2h1j | angiotensin converting enzyme | 2 | 607 | Crystal Structure of Drosophila AnCE | *Drosophila melanogaster* | 5,262 | 4,154 | 5,87 | 7,148 |
| 2h1n | Oligoendopeptidase F | 2 | 567 | 3.0 A X-ray structure of putative oligoendopeptidase F: crystals grown by vapor diffusion technique | *Geobacillus stearothermophilus* | 5,298 | 4,127 | 6,935 | 6,569 |

| | | | | | | | | | |
|---|---|---|---|---|---|---|---|---|---|
| 1y7910 | Peptidyl-Dipeptidase Dcp | 1 | 680 | Crystal Structure of the *E.coli* Dipeptidyl Carboxypeptidase Dcp in Complex with a Peptidic Inhibitor | *Escherichia coli* | 5,305 | 4,516 | 3,971 | 3,405 |
| 5l44 | K-26 dipeptidyl carboxypeptidase | 2 | 683 | Structure of K-26-DCP in complex with the K-26 tripeptide | *Astrosporangium hypotensionis K-26* | 5,367 | 3,99 | 3,444 | 2,971 |
| 5l43 | K-26 dipeptidyl carboxypeptidase | 2 | 683 | Structure of K26-DCP | *Astrosporangium hypotensionis K-26* | 5,405 | 4,326 | 3,541 | 3,085 |
| 2qr4 | Peptidase M3B, oligoendopeptidase F | 2 | 587 | Crystal structure of oligoendopeptidase-F from Enterococcus faecium | *Enterococcus faecium DO* | 5,535 | 5,217 | 6,642 | 6,973 |
| 3hq2 | Bacillus subtilis M32 carboxypeptidase | 2 | 501 | BsuCP Crystal Structure | *Bacillus subtilis* | 5,553 | 5,42 | 6,759 | 5,05 |
| 4ka7 | Oligopeptidase A, short endogenous peptide substrate | 1 | 714 | Structure of Organellar OligoPeptidase (E572Q) in complex with an endogenous substrate | *Arabidopsis thaliana* | 5,607 | 3,89 | 4,041 | 3,041 |
| 3ahn | Oligopeptidase | 2 | 564 | PZ PEPTIDASE A with Inhibitor 1 | *Geobacillus sp. MO-1* | 5,649 | 4,163 | 7,044 | 5,565 |

| | | | | | | | | | |
|---|---|---|---|---|---|---|---|---|---|
| 2o36 | Thimet oligopeptidase | 1 | 674 | Crystal structure of engineered thimet oligopeptidase with neurolysin specificity in neurotensin cleavage site | *Homo sapiens* | 4,05 | 5,497 | 0,124 | 1,049 |
| 3ahm | Oligopeptidase | 2 | 564 | Pz peptidase a | *Geobacillus sp. MO-1* | 5,675 | 3,942 | 6,627 | 5,405 |
| 5giv | Carboxypeptidase 1 | 6 | 503 | Crystal structure of M32 carboxypeptidase from Deinococcus radiodurans R1 | *Deinococcus radiodurans R1* | 5,765 | 5,234 | 5,145 | 5,463 |
| 3sks | Putative Oligoendopeptidase F | 1 | 567 | Crystal structure of a putative oligoendopeptidase F from Bacillus anthracis str. Ames | *Bacillus anthracis str. Ames* | 6,652 | 4,35 | 6,83 | 7,106 |
| 4fxyP | Neurolysin, mitochondrial | 2 | 693 | Crystal structure of rat neurolysin with bound pyrazolidin inhibitor | *Rattus norvegicus* | 4 | 4,98 | 1,236 | 0,486 |
