## Supplementary material for "A modular molecular framework for quickly estimating the binding affinity of the spike protein of SARS-CoV-2 variants for ACE2, in presence of mutations at the spike receptor binding domain": Supp. Tab. 3

| **Supp. Tab. 3 *In vitro* binding assays for the estimation of the binding afinity between the investigated SARS-CoV-2 spike RBD mutants and ACE2, compared to the binding affinity calculated for the Wuhan SARS-CoV-2 spike RBD/ACE2** | | | | | |
|---|---|---|---|---|---|
| **Single mutants** | **Tian F, Tong B, Sun L, et al (2021) Mutation N501Y in RBD of Spike Protein Strengthens the Inter-action between COVID-19 and its Receptor ACE2. bioRxiv https://www.biorxiv.org/content/10.1101/2021.02.14.431117v2** | | **Lopez E, Haycroft ER, Adair A, et al (2021) Simultaneous evaluation of antibodies that inhibit SARS-CoV-2 RBD variants with a novel competitive multiplex assay. medRxiv doi.org/10.1101/2021.03.20.21254037** | | **Deshpande A, Harris BD, Martinez-Sobrido L, et al (2021) Epitope classification and RBD binding properties of neutralizing antibodies against SARS-CoV-2 variants of concern. bioRxiv 2021.04.13.439681. https://doi.org/10.1101/2021.04.13.439681** |

| SARS-CoV-2 spike RBD Single Mutants/ACE2 affinity estimations | FACS ACE2 (The full-length ACE2 construct contains the ACE2 protein (residues 1-805), followed by a GGSGGGGS linker) cells, RDB 5 µM to 0.25 nM with 3-fold dilution | SPR Concentrations used for ACE2 protein were 50, 20, 10, 5, 2, and 1 nM respectively. Values were fitted to the 1:1 binding model. | AFM-SMFS the strength between RBDs and ACE2 on the living cell | | BLI profiles Global fit 1:1 30°C- 3µg/ml of biotinylated ACE2 (truncated human ACE2 ectodomain (residues 19–613) with a C495 -terminal AVI-tag and 6xHis-tag) ligand for 180s- 2-fold serial dilutions of each recombinant SARS-CoV-2 RBD variant | RDB multiplex assay 20µl of RBD natural variant cocktail - 10 µl of mAbs (final concentration 80nM per well) 8-point 4-fold titrations ACE2 inhibition assay 20µl of 25µg/ml of AviTagged Biotinylated ACE2 volume per well to 50µl. Incubation time 2 hours RT | | SPR Soluble ACE2 was injected over the RBD surfaces at 5 concentrations (300 nM, 75 nM, 18.75 nM, 4.7nM, and 1.2 nM) at a flow rate of 40 µL/min. The resulting sensorgrams were globally fit to a 1:1 model |
|---|---|---|---|---|---|---|---|---|
| spike RBD amino acid replacement | KD (nM) | KD (nM) | force pN | | KD (nM) | EC50 (µg/ml) | | KD (nM) |
| N439K | | | | | 25,2 | 12,2 | | |
| E484K | 38,5 ± 12 | | | | 37,2 | 23,8 | | |

| | | | | | | | | |
|---|---|---|---|---|---|---|---|---|
| **Wuhan SARS-CoV-2 Spike RBD/ACE2 (6m0j.pdb)** | 56,9 ± 16 | 8,3 ± 8,1 | 49 ± 11 | | 29,4 | 13,5 | | 80 |
| **L452R** | | | | | | | | 34,4 |
| **N501Y** | 13,1 ± 3,8 | 0,5 ± 0,5 | 57 ± 18 | | 6,4 | 0,2 | | 72,5 |
| **K417N** | 145 ± 45 | | | | 26,1 | | | |
| **S494P** | | | | | | 6,3 | | |
| **E484Q** | | | | | | 20,5 | | |
| **S477N** | | | | | 10 | 8,3 | | |
| **K417T** | | | | | | | | 131 |

**Supp. Tab. 3 *In vitro* binding assays for the estimation of the binding afinity between the investigated SARS-CoV-2 spike RBD mutants and ACE2, compared to the binding affinity calculated for the Wuhan SARS-CoV-2 spike RBD/ACE2**

| | | | | | |
|---|---|---|---|---|---|
| **Double/Triple Mutants** | **Tian F, Tong B, Sun L, et al (2021) Mutation N501Y in RBD of Spike Protein Strengthens the Inter-action between COVID-19 and its Receptor ACE2. bioRxiv https://www.biorxiv.org/content/10.1101/2021.02.14.431117v2** | | | | **Lopez E, Haycroft ER, Adair A, et al (2021) Simultaneous evaluation of antibodies that inhibit SARS-CoV-2 RBD variants with a novel competitive multiplex assay. medRxiv doi.org/10.1101/2021.03.20.21254037** |
| **SARS-CoV-2 spike RBD Single Mutants/ACE2 affinity estimations** | **FACS ACE2 (The full-length ACE2 construct contains the ACE2 protein (residues 1-805), followed by a GGSGGGGS linker) cells, RDB 5 μM to 0.25 nM with 3-fold dilution** | **SPR Concentrations used for ACE2 protein were 50, 20, 10, 5, 2, and 1 nM respectively. Values were fitted to the 1:1 binding model.** | **AFM-SMFS the strength between RBDs and ACE2 on the living cell** | | **microscale thermophoresis Recombinant ACE2 (full lenght) recombinant RBD; ratio 1:1 of ACE2 dilution and RBD labeled (ratio 2:1 protein: dye)** |
| **SARS-CoV-2 spike RBD amino acid replacement** | **KD (nM)** | **KD (nM)** | **force (pN)** | | **Kd (nM)** |
| **B.1.315_S.africa N501Y_E484K_K417T** | | | | | 87.6±25.5 |
| **Wuhan SARS-CoV-2 Spike RBD/ACE2 (6m0j.pdb)** | 56,9 ± 16 | 8,3 ± 8,1 | 49 ± 11 | | 402.5±112.1 |

| | | | | | |
|---|---|---|---|---|---|
| **P1_Japan/Brazil N501Y_E484K_K417N** | 45,2 ± 13,9 | 0,5 ± 0,5 | 56 ± 12 | | |
| **B.1.1.7_UK S494P_N501Y_E484K** | | | | | 203.7±57.1 |
